# Spatiotemporal development of the human T follicular helper cell response to Influenza vaccination

**DOI:** 10.1101/2023.08.29.555186

**Authors:** Stefan A Schattgen, Jackson S. Turner, Mohamed A Ghonim, Jeremy Chase Crawford, Aaron J. Schmitz, Hyunjin Kim, Julian Q. Zhou, Walid Awad, Wooseob Kim, Katherine M. McIntire, Alem Haile, Michael K. Klebert, Teresa Suessen, William D. Middleton, Sharlene A. Teefey, Rachel M. Presti, Ali H. Ellebedy, Paul G. Thomas

## Abstract

We profiled blood and draining lymph node (LN) samples from human volunteers after influenza vaccination over two years to define evolution in the T follicular helper cell (TFH) response. We show LN TFH cells expanded in a clonal-manner during the first two weeks after vaccination and persisted within the LN for up to six months. LN and circulating TFH (cTFH) clonotypes overlapped but had distinct kinetics. LN TFH cell phenotypes were heterogeneous and mutable, first differentiating into pre-TFH during the month after vaccination before maturing into GC and IL-10+ TFH cells. TFH expansion, upregulation of glucose metabolism, and redifferentiation into GC TFH cells occurred with faster kinetics after re-vaccination in the second year. We identified several influenza-specific TFH clonal lineages, including multiple responses targeting internal influenza proteins, and show each TFH state is attainable within a lineage. This study demonstrates that human TFH cells form a durable and dynamic multi-tissue network.

## Introduction

Neutralizing humoral immune responses against Influenza virus (Flu) are a key correlate of protection and the goal of vaccination. The generation of high-affinity antibodies takes place in the germinal centers (GCs) located in secondary lymphoid organs (SLO) where they are selected through affinity-maturation. CD4+ T follicular helper (TFH) cells provide support to B cells in the GC reaction through the production of cytokines and costimulatory molecule signaling engaged through T-B cell contact ^1^. TFH cells expressing the chemokine receptor CXCR5 and programmed death 1 (PD-1, gene name *PDCD1*) are readily detectable in SLOs and tertiary lymphoid organs supporting GC formation. While the frequency of circulating TFH (cTFH) in the peripheral blood is typically low in the absence of infection or vaccination ^2,3^, cTFH expansion and activation in response to vaccination correlated positively with the generation of high-affinity, class-switched antibodies ^4^. Recent evidence shows a diverse polyfunctional T cell response, including those cTFH cells, is correlated with protection against symptomatic infection independent of serology ^5^. Considering their influence on the quality of humoral immune response, a better understanding of the mechanisms regulating the TFH response to vaccination is necessary for designing improved vaccine platforms.

TFH cells display substantial heterogeneity and mutability in their phenotypes across location and time, and each of these states is not equal in their ability to boost and support B cell responses. For example, the Th1-like CXCR3^+^ CXCR5^low^ PD-1^low^ cTFH subset has impaired capacity for B cell help compared to CXCR3^-^ cTFH cells which more closely resemble GC TFH cells ^6–8^. As cTFH cells are replenished by GC TFH cells egressing from SLOs, TFH cells in these two compartments display significant overlap in their core gene expression (GEX) programs with tissue-specific modifications between lymphoid organs and blood ^9,10^. Antigen-specific TFH cells generated in response to vaccination or infection are able to form long-lived memory responses and be later recalled to assist B cells in subsequent challenges ^3,11^. In response to Flu vaccination in particular, the increased frequency of cTFH cells correlates with the induction of protective antibody responses ^4,11,12^. Deep repertoire profiling of tonsillar and blood TFH cells for matched donors showed a high frequency of shared clonotypes between the two compartments and found Flu-specific clonotypes shared across tissues ^10,13^.

Studies profiling the GEX and TCR repertoires of bulk TFH cell populations have revealed that they operate in a multi-tissue network linked by their TCR specificity. However, the trajectory of TFH differentiation and maturation simultaneously within SLOs and peripheral blood of humans in response to vaccination has not been tracked over time. Previously, we described the clonal evolution of the GC B cell response following Flu vaccination by serially-sampling lymph nodes via fine needle aspirate (FNA) and blood of human volunteers and performing single-cell GEX and immune repertoire sequencing ^14^. Here, we focused on characterizing the qualities and kinetics of the LN and cTFH response accompanying robust vaccine-induced GC reactions in multiple donors over the course of two years.

We found that alongside a transient increase in the frequency of cTFH cells in the blood during the first two weeks post-vaccination, LN TFH cells had expanded clonally and began differentiating into interfollicular pre-TFH cells before slowly maturing into GC TFH over the course of one to three months. In addition to the classical GC TFH cells, we observed the emergence of the recently described IL-10+ TFH cell subset ^15,16^ by two months post-vaccination, characterized by a unique transcriptional profile compared to pre-TFH and GC TFH subsets. Closer examination of the transcriptional profiles of TFH cells over time showed the existence of distinct phenotypic states along the trajectory of differentiation, and that each of these states is obtainable and dynamic within individual clonal lineages. Importantly, the key states and transitions occurring in the LN were not mirrored in the cTFH compartment, which had dramatically different kinetics. Together, these data present a spatio-temporal view of the dynamic changes in the repertoires and phenotypes of TFH cells engaged after successful vaccination and help define a standard regarding the desired TFH response in optimizing vaccine design.

## Results

### Activation of the TFH response following vaccination

Our cohort contains five healthy young adult volunteers enrolled in a multi-year seasonal Flu vaccination study (details in Methods). These donors had not received a seasonal Flu vaccine for at least three years prior to enrollment. Each donor was administered the Northern Hemisphere seasonal quadrivalent Flu vaccine (2018-2019 for year 1) and blood and/or LN samples were collected before vaccination, at 1 and 2 weeks and at approximately 1, 2, 3, 4, and 6 months after vaccination. Donors 321-05 and 321-04 were vaccinated the following year with the 2019-2020 seasonal quadrivalent vaccine and additionally sampled at 1 and 2 weeks and at 1, 2, 3, and 4 months after their second vaccination. Donor 321-05 was the most comprehensively sampled with corresponding blood and LN biopsies collected at nearly all time points during year 1, and LN samples at five additional time points after revaccination during year 2 (**Fig 1A**). Flow cytometric analysis of the year 1 time points for 321-05 showed that the frequency of CD4^+^CXCR5^+^CD38^+^ cTFH cells in the blood was low at 0.16% of CD4+ T cells prior to vaccination, expanded by about four-fold to their peak at 5 days afterward before contracting to baseline levels at day 28 (**Fig 1B-D**, Gating strategies for PBMC and LN TFH can be found in **S1A-B)**. A similar transient increase in cTFH frequency early after vaccination during year 1 was seen in donor 321-04 **(Fig S1C)**. Day 5 cTFH cells had increased levels of the activation markers ICOS and CD71 compared to the other time points (**Fig 1C**). The frequency of CXCR5^+^PD-1^hi^ TFH cells in the lymph nodes began increasing at day 5 and peaked at day 60 before contracting at the later time points (**Fig 1E**), and expressed high levels of the master transcription factor for TFH development B-Cell Lymphoma 6 protein (BCL-6) (**Fig 1D, F)**. The frequency of TFH cells positively correlated with those of GC B cells in the LN (**Fig 1D + S1D)**.

**Figure 1.**
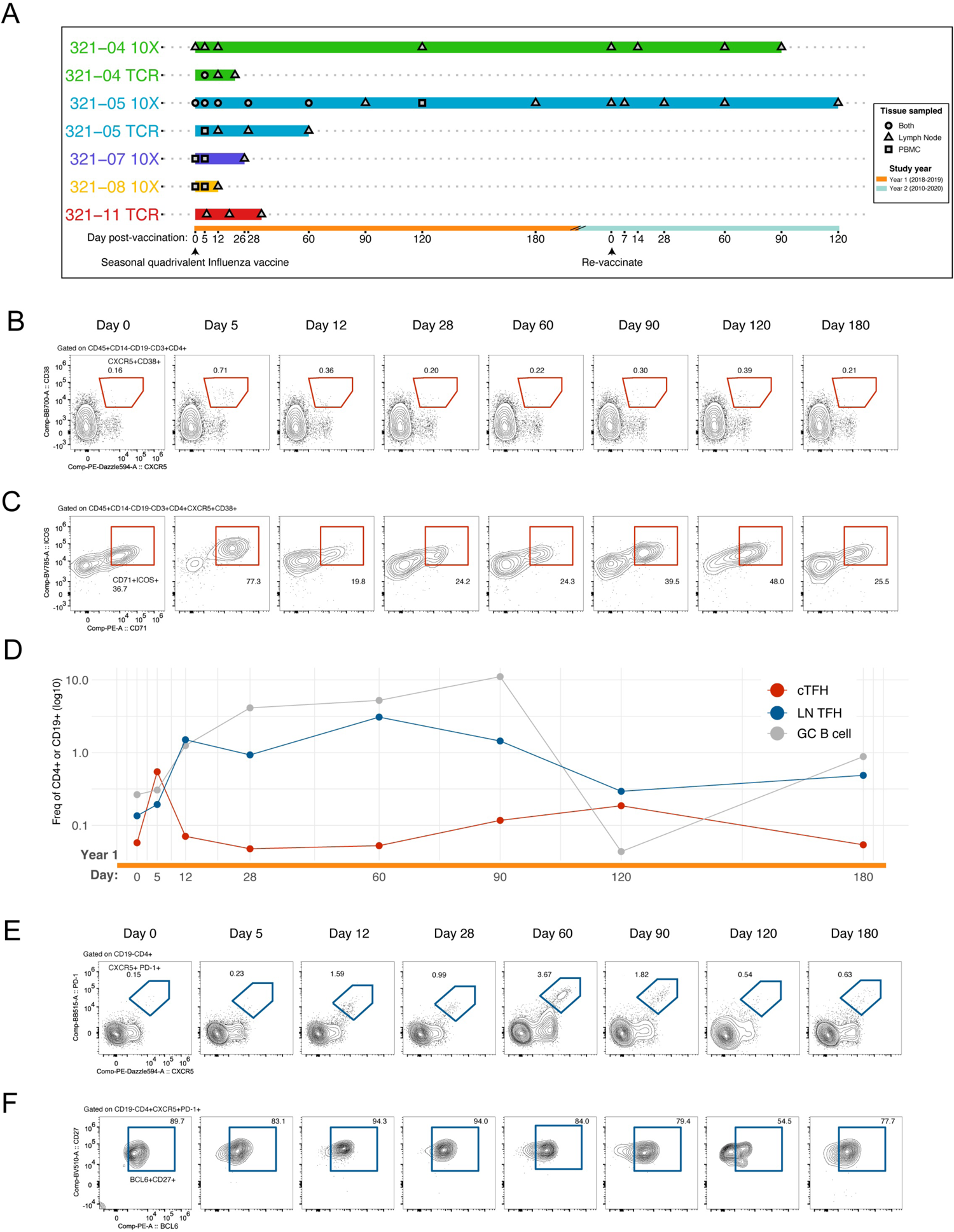
Detection of human TFH response in multiple tissues following seasonal Flu vaccination. **A)** Schematic of PBMC draws and lymph node fine needle aspirates (LN) for the three donors following seasonal influenza vaccination. **B)** Scatter plots of cTFH (CXCR5+CD38+) cells in the blood for donor 321-05 during year 1. Gated on CD45+ CD14-CD19-CD3+ CD4+ cells. See figures S1A-B for gating strategy. **C)** Frequency of ICOS+ CD71+ cTFH cells gated in B. **D)** Frequency of PBMC cTFH and LN TFH (CD4+ CXCR5+ PD-1+ BCL6+) and GC B cells (CD19+ IgDlo CD20hi CD38int) by time point. Shown as log10 frequency of CD4+ T cells or CD19+ B cells. **E)** Scatter plots of TFH (CXCR5+PD-1+) cells in LN samples. Gated on CD14-CD19-CD4+ cells. **F)** Frequency of lymph node CD27+ BCL6+ TFH cells gated in E.

To characterize changes in T cell phenotype elicited by repeated Flu vaccination, we performed single-cell gene expression (scGEX) and TCR sequencing on all immune cell types present in either peripheral blood or the same lymph node serially sampled from donors 321-04 and 321-05 over the course of two years, and lymph node samples days 12 and 26 during the first year for donors 321-08 and 321-07, respectively (32 unique tissue/time/donor combinations from 56 individual libraries) (**Table 1**). First, the gene count matrices for all the samples from both donors were aggregated into a single matrix containing all cell types and TCR data mapped to their corresponding cells. Next, we subset on cells belonging to clusters expressing T cell markers and paired TCR clonotype information to create a dataset containing exclusively T cells that included 154.547 cells covered by 127,471 unique TCR clonotypes (See Methods for more details on data processing). Each cluster could be broadly categorized as being naive, effector, or memory CD4+ and CD8+ T cells (*CCR7, SELL, KLRG1, ICOS, FOXP3, CD8B,* and *CD4*), mucosal-associated invariant T cells (MAIT) and NK T cells (*KLRB1* and TCR sequence), or TFH cells (*CXCR5 and PDCD1*) based on the expression of marker genes indicative for each subset (**Fig 2A, 2C, 2G, and S2A)**. Closer inspection of the factors associated with each T cell cluster showed that donor, tissue source, and study year all were significant factors influencing T cell GEX (**Fig 2B, 2D-F, S2B-E)**. The degree of clonal expansion within each cluster was consistent with their naive and effector/memory GEX profiles; naive clusters had no clones with more than 2 cells whereas effector/memory, MAIT/NKT, and TFH clusters contained a number of extensively expanded clones **(Fig S2F-G)**. One cluster corresponding to TFH cells showed higher and specific expression of characteristic markers including *CXCR5*, *PDCD1*, *ICOS*, *IL21*, and *TOX2,* and contained cells from all four donors (**Fig 2G**).

**Figure 2.**
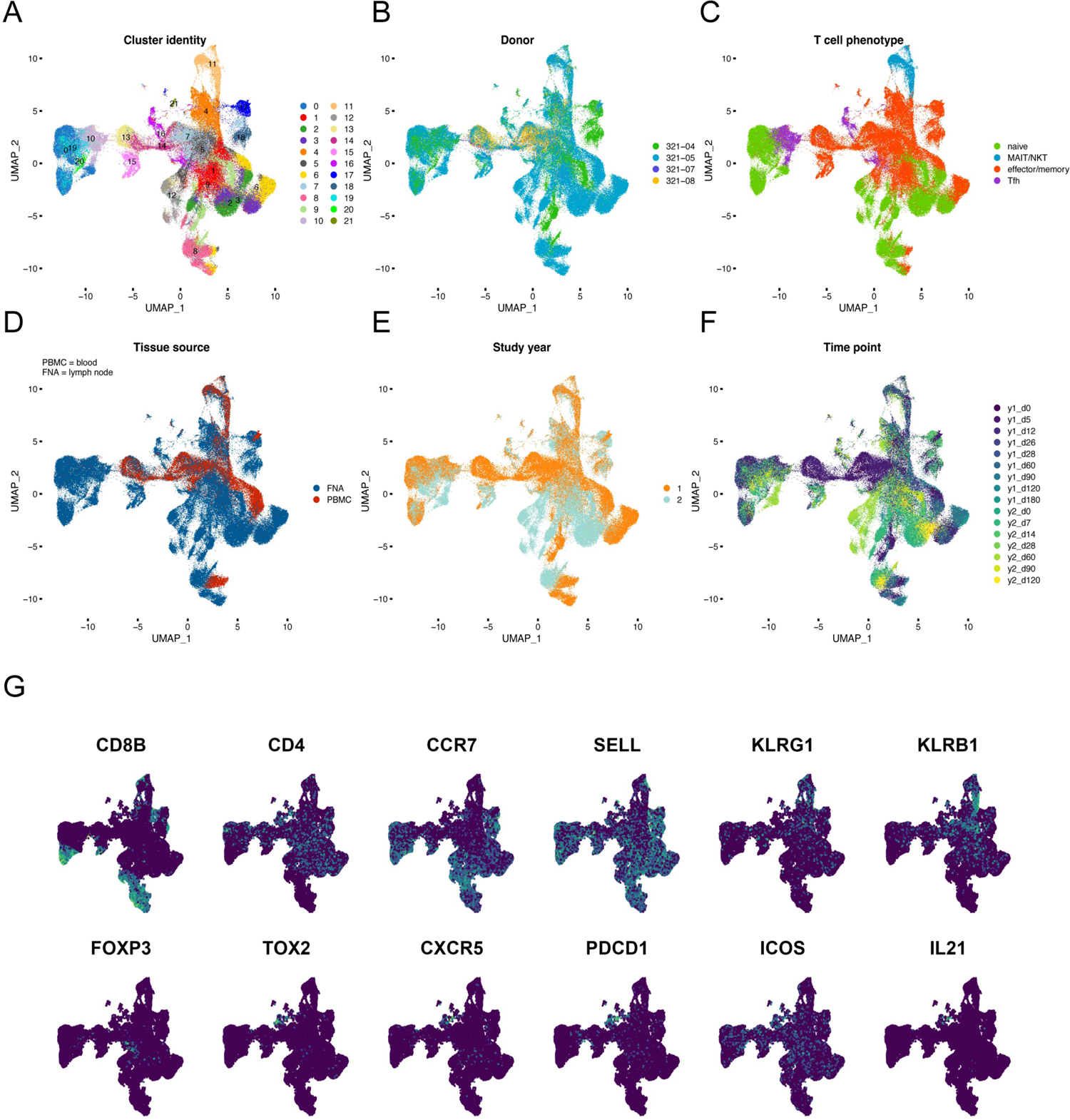
Spatial-temporal GEX and TCR repertoire profiling of T cell response to Flu vaccination. Aggregate scGEX dataset containing N = 154,547 cells with 127,471 unique TCR clonotypes generated from PBMC and LN samples at each time point for donors 321-05 and 321-04 (See Figure 1A). 2D umap projection annotated by: **A)** GEX cluster T cell phenotype, **B)** donor origin, **C)** T cell type, **D)** tissue origin, **E)** study year, and **F)** clone size. **G)** Feature plots displaying select T subset markers.

**Table 1.** Cell and clonotype counts in total T cell dataset.

| donor | year | day | tissue | number of cells | number of clonotypes |
| --- | --- | --- | --- | --- | --- |
| 321-04 | 1 | 0 | LN | 4685 | 4430 |
| 321-04 | 1 | 12 | LN | 4593 | 4443 |
| 321-04 | 1 | 120 | LN | 2686 | 2426 |
| 321-04 | 1 | 5 | LN | 1717 | 1642 |
| 321-04 | 2 | 0 | LN | 4007 | 3859 |
| 321-04 | 2 | 14 | LN | 5212 | 5035 |
| 321-04 | 2 | 60 | LN | 1546 | 1470 |
| 321-04 | 2 | 90 | LN | 2711 | 2555 |
| 321-05 | 1 | 0 | LN | 6613 | 6426 |
| 321-05 | 1 | 12 | LN | 8859 | 8493 |
| 321-05 | 1 | 180 | LN | 7199 | 7003 |
| 321-05 | 1 | 28 | LN | 6794 | 6507 |
| 321-05 | 1 | 5 | LN | 7311 | 7109 |
| 321-05 | 1 | 60 | LN | 4336 | 4133 |
| 321-05 | 1 | 90 | LN | 6083 | 5888 |
| 321-05 | 2 | 0 | LN | 11245 | 10979 |
| 321-05 | 2 | 120 | LN | 4670 | 4531 |
| 321-05 | 2 | 28 | LN | 5279 | 5117 |
| 321-05 | 2 | 60 | LN | 6234 | 5946 |
| 321-05 | 2 | 7 | LN | 12798 | 12509 |
| 321-05 | 1 | 0 | PBMC | 3353 | 2778 |
| 321-05 | 1 | 12 | PBMC | 3584 | 3136 |
| 321-05 | 1 | 120 | PBMC | 2828 | 2379 |
| 321-05 | 1 | 28 | PBMC | 3121 | 2717 |
| 321-05 | 1 | 5 | PBMC | 8730 | 5220 |
| 321-05 | 1 | 60 | PBMC | 3611 | 3027 |
| 321-07 | 1 | 26 | LN | 722 | 412 |
| 321-07 | 1 | 0 | PBMC | 944 | 559 |
| 321-07 | 1 | 5 | PBMC | 1632 | 953 |
| 321-07 | 1 | NA | PBMC | 130 | 41 |
| 321-08 | 1 | 12 | LN | 6961 | 3057 |
| 321-08 | 1 | 0 | PBMC | 2554 | 1406 |
| 321-08 | 1 | 5 | PBMC | 1491 | 768 |
| 321-08 | 1 | NA | PBMC | 308 | 133 |

### TFH cells exist in distinct transcriptional states along the trajectory of maturation

We next focused on the TFH cells to characterize their phenotypic plasticity over time in response to Flu vaccination. CD4 helper T cells can differentiate into multiple subsets within clonal lineages stemming from a single founder. With this in mind, we subset all cells in the entire dataset with TCR sequences matching those cells contained within the TFH cluster.

Interestingly, regulatory CD4+ T cells (Tregs) were located near to the TFH cells within the embedding indicating they shared some transcriptional features perhaps due to localization within SLOs near GC structures. Tregs have also been shown to regulate GC responses in mouse models and prompted us to hypothesize they play a similar role in humans ^17–19^. This strategy to capture TFH and Treg cells yielded 15,290 cells with 11,268 unique TCR clonotypes (**Table 2**). Reclustering of the subset discovered 20 GEX clusters where again the donor of the cells accounted for much of the variation in the GEX (**Fig 3A-B**). These cells were predominantly from the LN samples; however, 1,710 cells were from the peripheral blood indicating our approach captured both the central and lymph node TFH populations and their clonal relations (**Fig 3C**). In addition to donor variation, the GEX varied substantially across sampling time points roughly partitioned into early (top), middle (bottom), and late (middle) time points in the projection (**Fig 3D**). Since the single-cell GEX dataset also contained lymph node B cells (see Methods for details on data processing), we looked for correlations between the frequency of the TFH cells and different B cell subsets in the lymph node. While there was no relationship between the frequencies of TFH cells and other B cell subsets, there was a significant positive correlation with GC B cell frequency (**Figs 3E, S3A)**.

**Figure 3.**
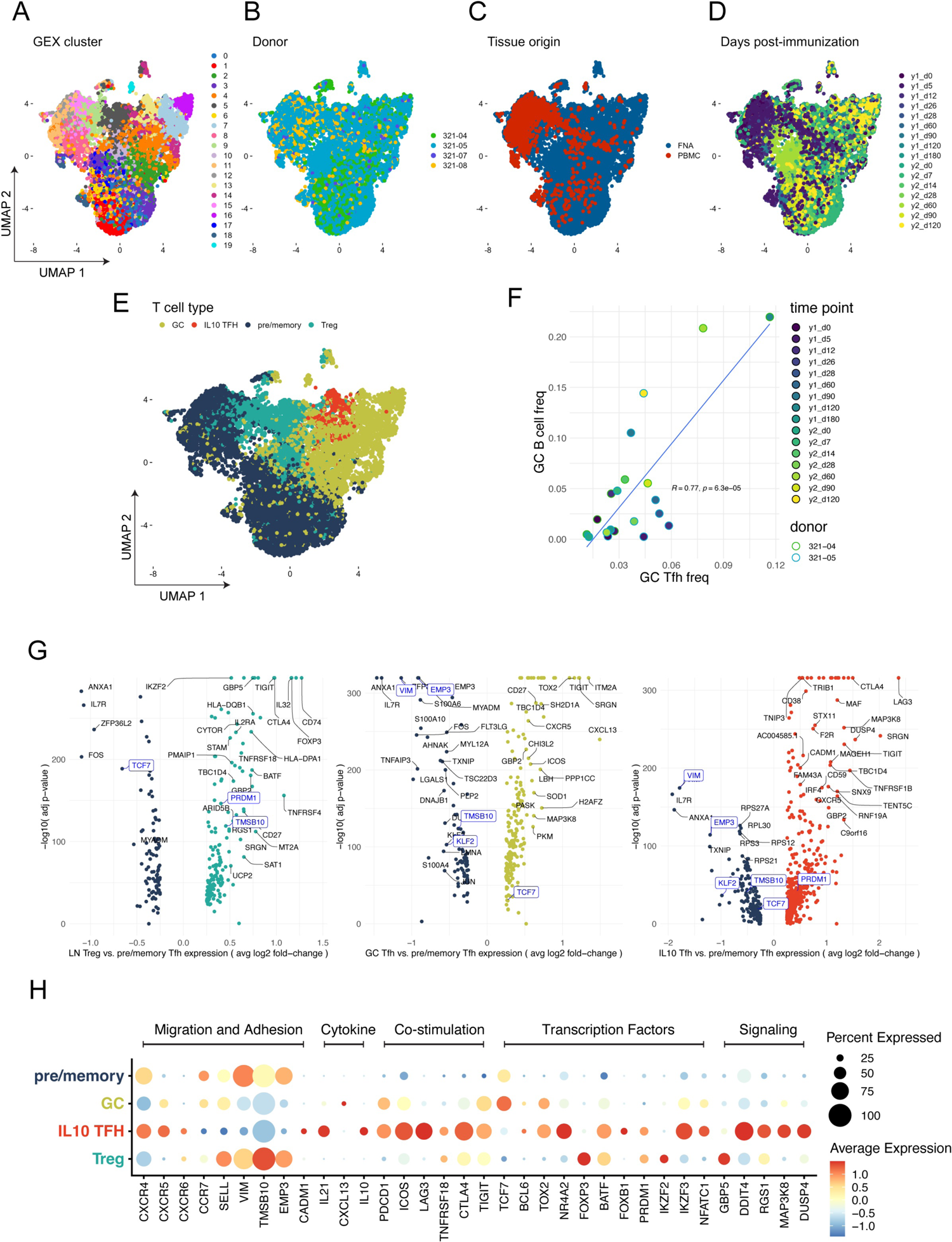
Identification of TFH maturation states. Subset of the aggregate T cell dataset in figure 2 containing all cells clonally related to those in the TFH and Treg clusters. N = 15,290 cells with 11,268 unique TCR clonotypes. 2D umap projections of the TFH lineage dataset annotated by: **A)** GEX cluster, **B)** donor, **C)** tissue, and **D)** time post-immunization. **E)** Scatter plot of comparing the relative frequencies of germinal center B cells versus TFH cells in LN samples. Pearson correlation. **F)** 2D umap projection of the TFH lineage dataset annotated by TFH subset. **G)** Volcano plots of TFH subset-specific marker genes for pre/memory vs. IL10 TFH (left), Treg (middle), and GC TFH (right). **H)** Dot plot of select TFH and Treg subset-specific markers.

**Table 2.** Cell and clonotype counts in TFH lineages dataset.

| donor | year | day | tissue | number of cells | number of clonotypes |
| --- | --- | --- | --- | --- | --- |
| 321-04 | 1 | 0 | LN | 252 | 245 |
| 321-04 | 1 | 5 | LN | 92 | 90 |
| 321-04 | 1 | 12 | LN | 324 | 287 |
| 321-04 | 1 | 120 | LN | 557 | 420 |
| 321-04 | 2 | 0 | LN | 264 | 263 |
| 321-04 | 2 | 14 | LN | 461 | 408 |
| 321-04 | 2 | 60 | LN | 255 | 226 |
| 321-04 | 2 | 90 | LN | 372 | 334 |
| 321-05 | 1 | 0 | LN | 627 | 611 |
| 321-05 | 1 | 5 | LN | 1646 | 1596 |
| 321-05 | 1 | 12 | LN | 1608 | 1415 |
| 321-05 | 1 | 28 | LN | 1443 | 1408 |
| 321-05 | 1 | 60 | LN | 677 | 654 |
| 321-05 | 1 | 90 | LN | 640 | 575 |
| 321-05 | 1 | 180 | LN | 744 | 705 |
| 321-05 | 2 | 0 | LN | 660 | 620 |
| 321-05 | 2 | 7 | LN | 931 | 819 |
| 321-05 | 2 | 28 | LN | 638 | 607 |
| 321-05 | 2 | 60 | LN | 583 | 565 |
| 321-05 | 2 | 120 | LN | 481 | 420 |
| 321-05 | 1 | 0 | PBMC | 140 | 127 |
| 321-05 | 1 | 5 | PBMC | 590 | 472 |
| 321-05 | 1 | 12 | PBMC | 180 | 167 |
| 321-05 | 1 | 28 | PBMC | 151 | 147 |
| 321-05 | 1 | 60 | PBMC | 149 | 141 |
| 321-05 | 1 | 120 | PBMC | 90 | 87 |
| 321-07 | 1 | 26 | LN | 53 | 53 |
| 321-07 | 1 | 0 | PBMC | 71 | 56 |
| 321-07 | 1 | 5 | PBMC | 27 | 27 |
| 321-07 | 1 | NA | PBMC | 2 | 2 |
| 321-08 | 1 | 12 | LN | 272 | 217 |
| 321-08 | 1 | 0 | PBMC | 197 | 159 |
| 321-08 | 1 | 5 | PBMC | 98 | 91 |
| 321-08 | 1 | NA | PBMC | 15 | 15 |

Examination of differentially-expressed genes across the TFH clusters showed several clusters (2, 4, 7, 14, 16, and 19) expressed high levels of genes characteristic of mature GC TFH cells (e.g. *CXCR5*, *PDCD1*, *IL21*), while other clusters (0, 1, 3, 6, 8, 11, 12, 15, 17, and 18) expressed genes consistent with the early and memory stages of TFH differentiation (e.g. *KLF2*, *IL7R*, *CCR7*) **(Fig S3B)**. Cluster 13 expressed marker genes (e.g., *IL10* and *LAG3*) most consistent with the recently described subset of T follicular cells producing IL-10 (IL10 TFH) ^15,16^. Clusters 5, 9, and 10 expressed the Treg master transcription factor Forkhead Box P3 (*FOXP3)* **(Fig S3B)**. We then classified each of the TFH clusters as either pre/memory TFH, GC TFH, IL10 TFH, or Treg cells based on these markers (**Fig 3F**), and defined the GEX profiles specific to each state. In addition to *FOXP3*, Tregs expressed other canonical markers of the lineage including Interleukin 2 receptor alpha subunit (*IL2RA*), Basic Leucine Zipper ATF-Like Transcription Factor (*BATF*), and PR/SET Domain 1 (*PRDM1*, aka *BLIMP1*) (**Fig 3G, left panel, 3H)**. Compared to early and memory TFH cells, GC TFH cells showed increased levels of the co-inhibitory receptor T cell immunoreceptor with Ig and ITIM domains (*TIGIT*), the transcription factor TOX high mobility group box family member 2 (*TOX2*) ^20^, and the canonical TFH chemokine *CXCL13* (**Fig 3G, middle pane, 3H)**. Consistent with their high motility along the T cell zone border, amongst the top expressed genes in the early and memory TFH cells were those associated with cell adhesion and cytoskeletal rearrangement including Vimentin (*VIM*), Epithelial membrane protein 3 (*EMP3*), and Thymosin beta-10 (*TMSB10*), as well as Kruppel Like Factor 2 (*KLF2)*, a negative regulator of TFH differentiation ^21^ (**Fig 3G-H**). In addition to IL-10, the IL10 TFH cells expressed several genes that distinguished the subset from the pre/memory subset (**Fig 3G right panel, Fig 3H**). Transcription Factor 7 (*TCF7*) promotes TFH differentiation by suppressing the expression of Th1-promoting *PRDM1* ^22^. Consistent with this mechanism, the pre/memory and GC TFH expressed high levels of *TCF7* and low amounts of *PRDM1*; in contrast, IL10 TFH cells expressed low levels of *TCF7* and high levels of *PRDM1* (**Fig 3G-H**). *FOXP3* was not detected in IL10 TFH cells, however, they specifically expressed another member of the Forkhead Box family, *FOXB1,* which has no described role in regulating

T cell differentiation and function. IL10 TFH cells expressed a number of genes encoding for proteins known to modify intracellular signaling pathways including the negative regulator of mTORC1 signaling DNA Damage Inducible Transcript 4 (*DDIT4*), Regulator Of G Protein Signaling 1 (*RGS1*) which regulates G-protein signaling T cell chemotaxis, and two regulators of MAPK signaling, Mitogen-Activated Protein Kinase Kinase Kinase 8 (*MAP3K8*) and Dual Specificity Phosphatase 4 (*DUSP4*) (**Fig 3H**). The transcriptional profiles within each TFH state were consistent between the two donors, indicating these discrete states along the trajectory of TFH differentiation following vaccination were common between them (**Fig S3C)**.

We asked whether particular biological processes in the Gene Ontology (GO) database were enriched within the TFH and Treg subsets and examined their variation across both time and T cell subset. We conducted gene set variation analysis (GSVA) with the top enriched GO terms using IL10 TFH marker genes relative to other TFH and Treg subsets. IL10 TFH were chosen as the representative subset since they shared many DEGs with GC TFH as well as a number of subset-specific DEGs; thus they are representative of mature TFH cells. The pathways found to be enriched within the IL10 TFH could be broadly categorized into those related to cellular metabolism, TCR signaling, and cytokine-mediated signaling (**Fig 4A).** A gradient across the pre/memory, GC, and IL10 TFH subsets was observed with all pathways being mostly highly represented in IL10 TFH, followed GC TFH, and lastly the pre/memory TFH cells. Many of the pathways enriched in the GC and IL10 TFH cells were similarly upregulated in Tregs. Pathways related to glycolysis were particularly upregulated in the IL10 and GC TFH subsets.

**Figure 4.**
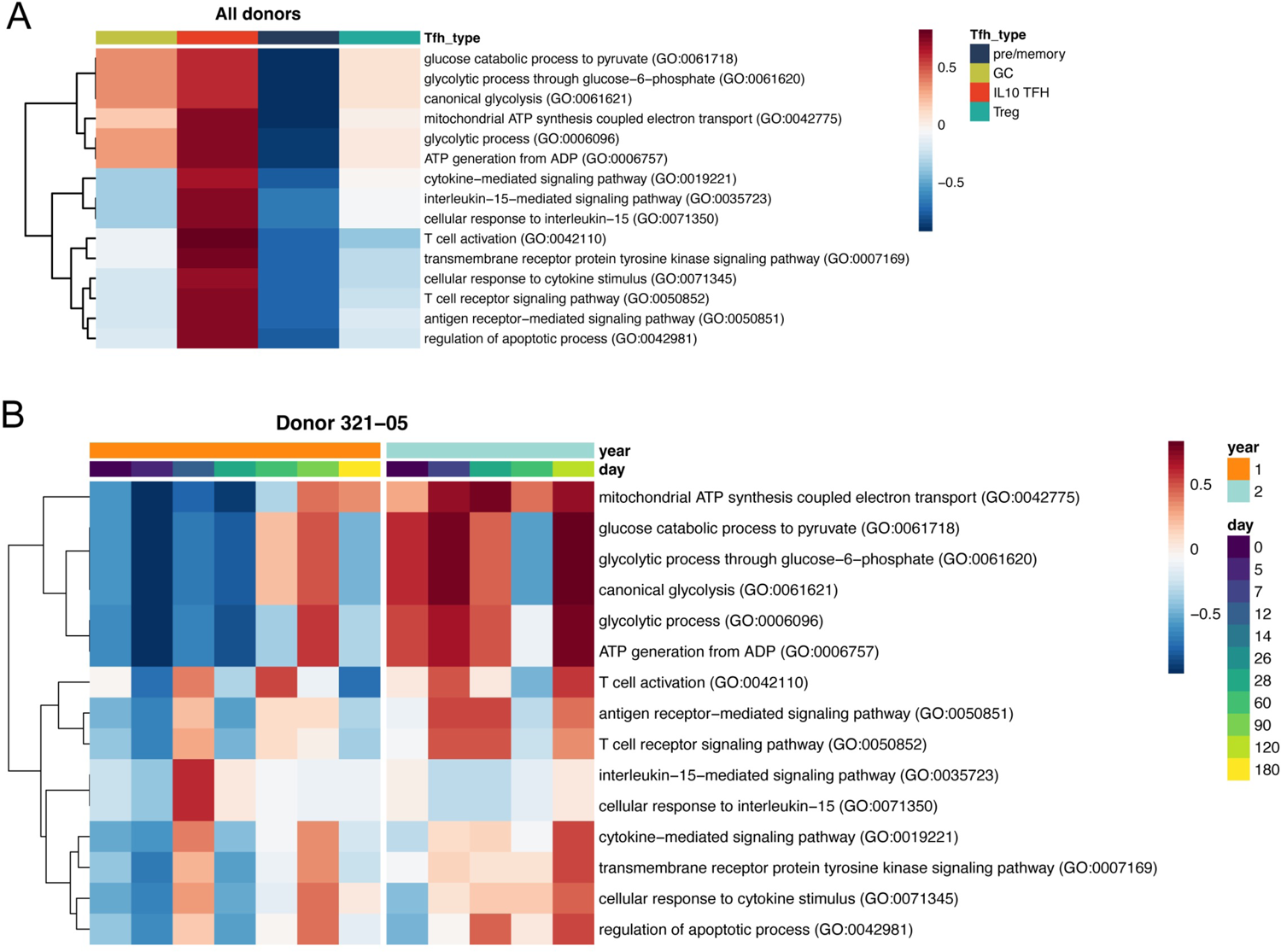
Alterations in TFH metabolism and signaling with time. **A)** GSVA of top upregulated GO terms identified by *Enrichr* analysis for each IL10 TFH subset relative to other TFH/Treg subsets. **B)** GSVA of donor 321-05 LN TFH cells (Tregs excluded) across time points..

GSVA was further performed on lymph node TFH cells for donors 321-05 and 321-04 separately, and in the absence of the Treg cells, to resolve changes in these pathways in TFH cells over the entire time course of the study. In donor 321-05 between days 0 and 28 of the first year, pathways associated with T cell activation and cytokine signaling were the most upregulated amongst all the pathways examined and peaked at day 60 (**Fig 4B**). Mitochondrial respiration and glycolysis pathways were sharply upregulated at days 60 and 90 before decreasing at day 180. In contrast to the slow evolution of the TFH response during year 1, the pathways describing T cell activation, cytokine-stimulation, and cellular metabolism were already higher at baseline on day 0 during year 2, and quickly increased by day 5 following revaccination. These pathways also varied in their expression across the time course for donor 321-04, however, in contrast to donor 321-05, glucose metabolism was upregulated by day 5 during the first year but displayed slower kinetics in upregulation the second year. **(Fig S4A)**. One potential interpretation is the TFH response to vaccination for donor 321-04 during year 1 was mostly recalled from memory, or a mix of primary and recall responses.

### Temporal dynamics of the TFH response

Considering the alterations in T cell activation and cellular metabolism pathways across TFH subsets, we next looked at changes in the relative frequency of each subset over time. In donor 321-05, the pre/memory TFH cells were the largest fraction (∼56%), followed by GC TFH cells (∼25%), Tregs (18%), and few IL10 TFH cells (>1%) at the time of first vaccination (**Fig 5A**).

**Figure 5.**
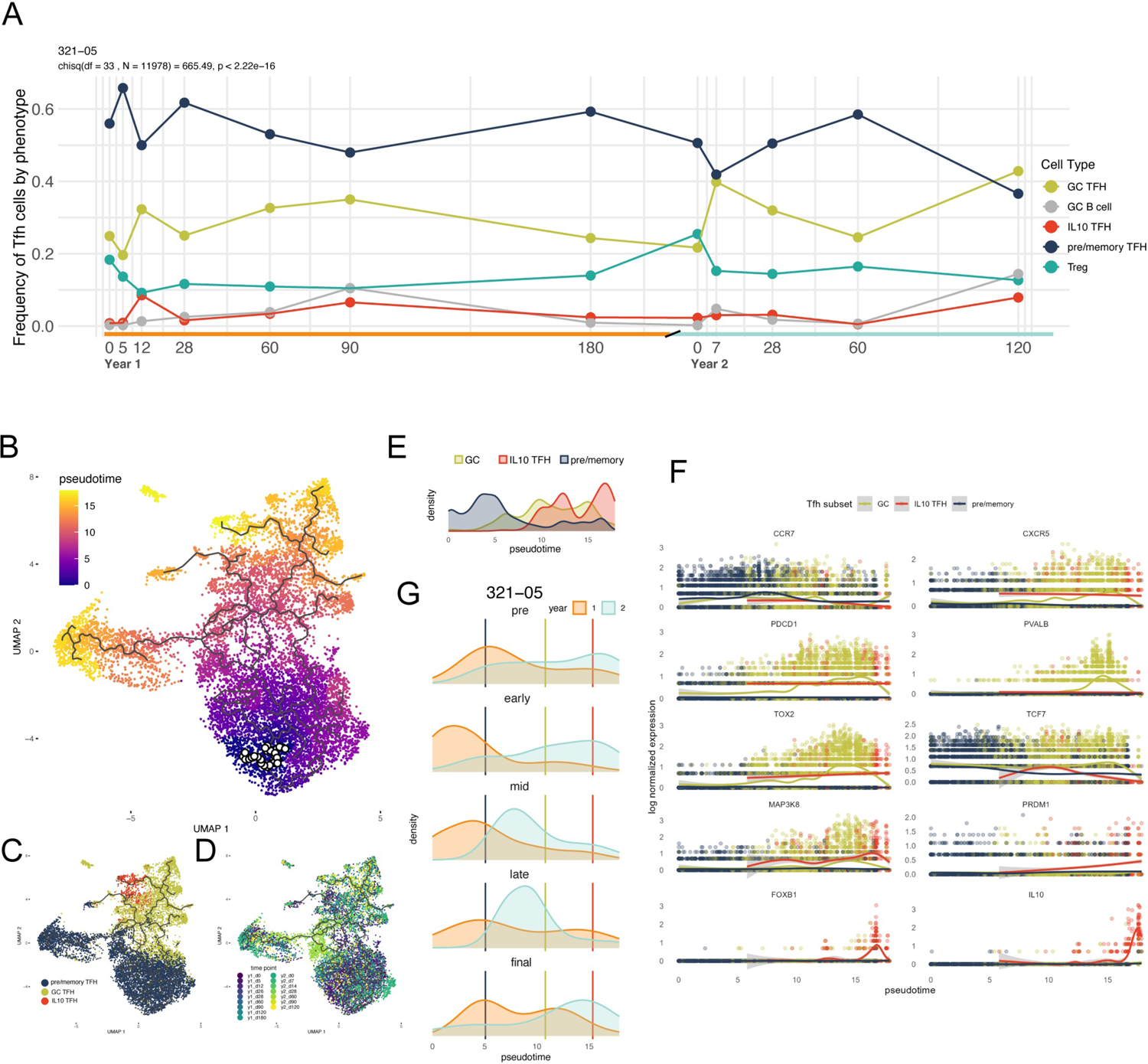
TFH composition and phenotype are dynamic. **A)** Relative frequencies of pre/memory, GC, and IL10 TFH subsets in LN samples from donor 321-05 over time. Chi-squared with p-value determined by Monte Carlo simulation. 2D umap projection of LN TFH cells (Tregs excluded) from all donors annotated by: **B)** pseudotime with selected root nodes where scale was pseudotime scale was set to zero indicated by black circles, **C)** TFH subset, and **D)** time point. **E)** Scatter plot of log normalized expression of TFH subset marker genes versuses pseudotime values per cell. **E)** Density plots of pseudotime values by TFH subset. **F)** Density plots of pseudotime values for 321-05 LN TFH cells for matched time points between study years. In the format (year 1 day, year 2 day), time points shown are pre (0,0), early (7,5), mid (28,28), late (60,60), final (180,120). Vertical lines indicate the median pseudotime from E) for the indicated TFH subset.

The frequency of GC TFH cells increased by day 12 and continued to rise until peaking at day 90 before declining to a similar frequency as day 0 at day 180. The frequency of IL10 TFH cells transiently increased at day 5 to ∼10% before declining and rising to ∼5% at day 90 at the peak of the GC B cell response. Tregs were highest in their frequencies at the earliest and latest time points and displayed an inverse pattern in their kinetics relative to the GC TFH cells. Pre/memory TFH cells were again the predominant subset at the time of re-vaccination on year 2 day 0. However, in contrast to year 1, the frequency of GC TFH had expanded two-fold by day 7 to 40% and remained in high frequency out to day 120. A corresponding two-fold decrease in Treg frequency was observed during this time like the inverse kinetics observed in year 1. IL10 TFH frequency increased during the first week after vaccination during year 2 and remained stable out through day 120. GC and pre/memory TFH cells were similar in frequency in Donor 321-04 on day 0 of year 1. Vaccination stimulated a transient increase in the frequency of GC IL10 TFH frequency by day 5 that had decreased to starting levels by day 12, whereas the frequency of Tregs decreased during this time and little change in GC B cell frequency was noted **(Fig S5A)**. The frequency of GC and IL10 TFH, and GC B cells, were all at their maximum at day 120. In contrast to year 1, the kinetics of the TFH response in donor 321-04 during year 2 showed a similar pattern as those observed in donor 321-05. Here, the frequency of the pre/memory TFH were highest at day 0 of year 2 and quickly declined as frequencies of GC and IL10 TFH, and GC B cells each increased and remained elevated through day 120.

We next performed pseudotime analysis to infer and quantify TFH maturation and compare their inferred developmental trajectories to those measured in real time. Here, we subset the dataset further into lymph node TFH cells only (excluding Tregs) and we set the “root” nodes (pseudotime values are set to 0) in an area of the projection containing mostly pre/memory TFH cells from early time points in year 1 (**Fig 5A-C).** Comparing the distributions of pseudotime values across TFH states showed the inferred pseudotime trajectories matched the expected our biological expectation; pre/memory subset were early in pseudotime while the more mature GC and IL10 TFH subsets were in middle and late pseudotime (**Fig 5E**). We next compared the expression of marker genes representative of the TFH subsets relative to pseudotime values to better assess the fit between the inferred and observed developmental trajectories. *CCR7* expression was highest in the early pseudotime pre/memory TFH cells whereas GC TFH markers *PDCD1 PVALB*, *TOX2*, and *CXCR5* increased in expression during mid to late pseudotime (**Fig 5F**). In line with our observations above, the *FOXB1* and *IL10* were specifically expressed by IL10 TFH cells in late pseudotime. Furthermore, *PRDM1* expression in late pseudotime IL10 TFH increased while expression of *TCF7* simultaneously decreased.

Given the changes in the kinetics of metabolism and signaling pathways between years 1 and 2, we compared the pseudotime distributions of matched time points across study years to better quantify differences in TFH maturation in the same lymph node across vaccination years. The mean pseudotime values in donor 321-05 steadily increased between early and final time points in year 1 and beyond, as the distribution skewed even further towards late pseudotime at day 0 in year 2 (**Fig 5G**). The pseudotime values across the year 2 time points were nearly all higher than those observed over the first year. Giving a contrasting view, the pseudotime values remain elevated across all the year 1 time points for donor 321-04 **(Fig S5B)**. However, the values dropped to a more expected baseline by the second year where the response now better matched that observed in donor 321-05 during year 1 where TFH maturation takes place over the course of weeks before fully maturing by 2-3 months. Together these observations show that TFH phenotypes are not only dynamically tuned in the short-term, but also over the long term to promote a robust and rapid memory recall response to antigen-restimulation.

### Clonal dynamics of the TFH cell response

We next turned our attention towards characterizing the TCR repertoire, specificity, and clonal dynamics of the TFH response. First, we asked if the frequency of TFH cells was related to their clonality. In donor 321-05, TFH frequency (amongst all T cells in the lymph node) during the first year increased between days 0 and 5, peaked at day 28 before slowly decreasing back to baseline at day 90 (**Fig 6A**). During year 1 in Donor 321-04, TFH frequency continued to increase across the entire time course and peaked at day 120. During the second year, TFH frequency quickly increased between days 0 and the first time point after revaccination (day 7 for 321-05, day 14 for 321-04) for both donors, and remained elevated through the final time points for the year.

**Figure 6.**
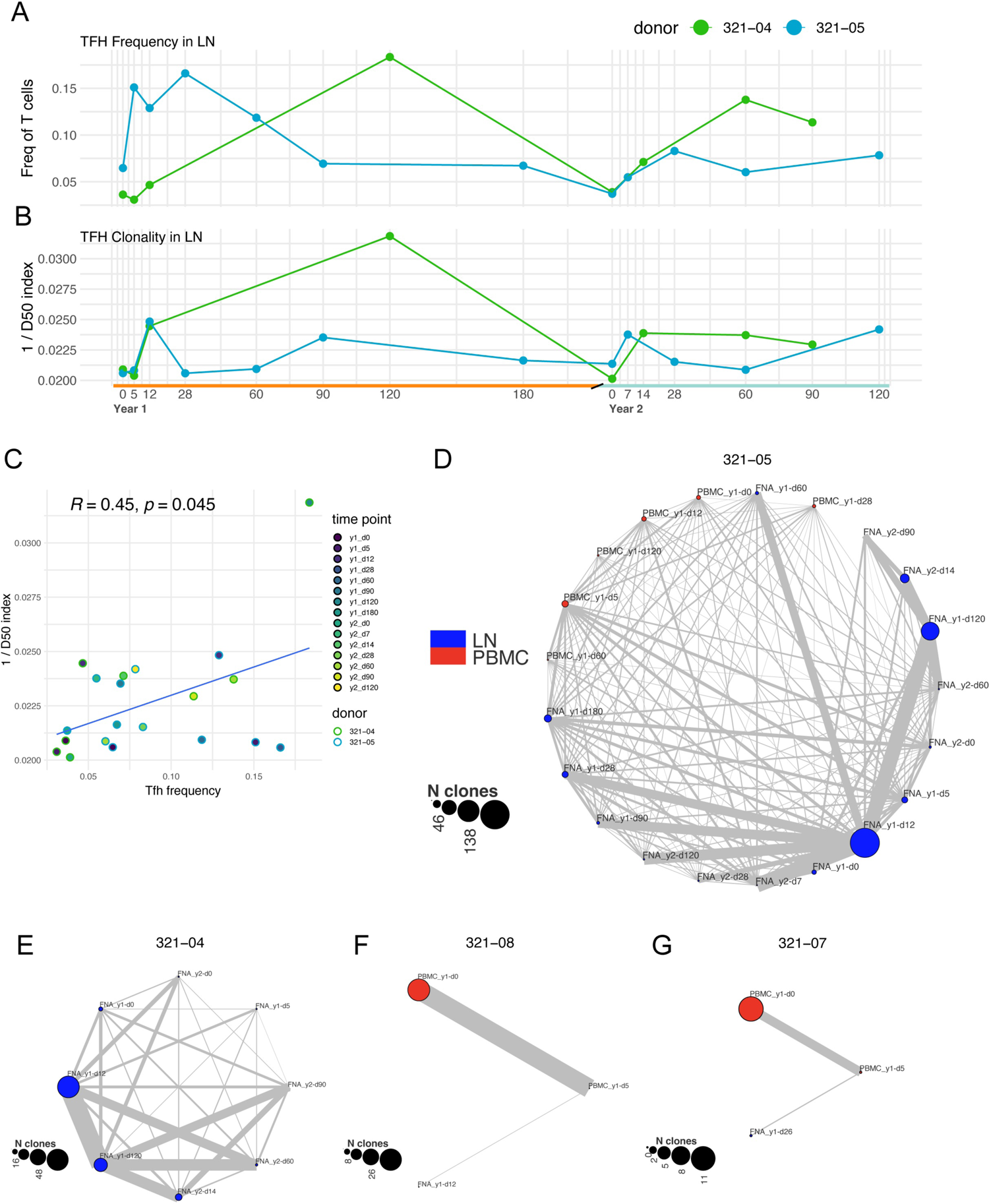
Clonal TFH expansion following flu vaccination. **A)** Relative frequency of TFH cells in LN samples with respect to time point. **B)** TFH clonality in LN samples with respect to time point as measured by inverse D50 index. **C)** Scatter plot of comparing the TFH relative frequency and clonality across all time points. Pearson correlation. **D-G)** Network graph depicting the connections between TFH clonal lineages in PBMCs and LN for donor 321-05 **D)**, 321-04 **E)**, 321-08 **F)**, and 321-07 **G)**. Node sizes correspond to the number of clones and edge widths correspond to the number of clones connecting each node.

We quantified the clonality of the TFH cells using the reciprocal of the D50 index (see methods). TFH clonality followed a similar trend to that observed for TFH frequency. During year 1, the peak in TFH frequency for donor 321-05 coincided with their maximum clonality at day 12 and a second peak without an accompanying spike in frequency was detected at day 90 (**Fig 6B**). For donor 321-04, the TFH clonality sharply increased at year 1 day 12, and remained elevated for the remainder of year 1. TFH frequency and clonality for both donors had contracted by the baseline time point at year 2 day 0, however, they quickly expanded in a clonal-manner by the first time point after revaccination and contracted over the next 2 months before again expanding and reaching a second peak at days 90 and 120 for donors 321-04 and 321-05, respectively. Comparing TFH frequency to their clonality showed a positive correlation between these two measures (**Fig 6C**), indicating TFH expansion occurs in a clonal-manner after vaccination.

Next, we tracked the TFH clonotypes across time points, and for donor 321-05 across tissues as well, to determine whether their repertoires remained stable over the long term. The number of TFH clonotypes in the lymph node remained low at year 1 days 0 and 5 prior to expanding on day 12 when the largest number of unique TFH clonotypes for donor 321-05 was observed (**Fig 6D**). Several of the TFH clones detected in the lymph node at year 1 day 12 were detected in peripheral blood at the year 1 day 0 time point indicating that at least some of the clonotypes engaging in the GC reaction were derived from memory cTFH cells. Additionally, many of the TFH clonotypes from year 1 day 12 were detected at subsequent time points, particularly at days 28, 60, 90, and 120, however, fewer were detectable by day 180. Very few TFH clonotypes persisted from year 1 until year 2 day 0, but by one week after revaccination (year 2 day 7) many of the clonotypes tracked from year 1 were again detectable and persisted throughout year 2 until the last time point at day 120. A similar trend was observed for donor 321-04 (**Fig 6E**). Here, several of the TFH clones detected at day 12 persisted throughout year 1, contracted by year 2 day 0, were detected again at year 2 day 14, and persisted through year 2 day 90. Shared clonotypes in the PBMC samples for donor 321-07 and 321-08 were detected across time points, and in both cases a single clonotype was shared between the blood and LN TFH compartments within the respective donor (**Fig 6F-G**).

To expand the clonal analysis and increase the depth of the repertoire data, we performed additional single-cell paired and bulk TCR sequencing on sorted cTFH and LN TFH cells from donors 321-04, 321-05, and 321-11 at multiple time points from year one (See **Fig 1A**). The additional paired TCR sequences from donors 321-04 and 321-05 matched mostly to cells within the TFH clusters in the 10X dataset **(Fig S6B)**. Matching TCR sequences obtained from bulk TCR profiling showed a similar pattern with most matches being to cells in the TFH clusters **(Fig S6C)**. We then used the Morisita-Horn overlap index to determine the extent of sharing, or publicity, between the bulk repertoire sequences from sorted cTFH and LN TFH cells. Here, sharing of TCR alpha and beta chains was highest amongst samples taken from the same donor, and consistent with observations from the 10X dataset, TCR sequences from blood cTFH on day 5 extensively overlapped with those from lymph node TFH cells at day 12 and later time points during year one for subjects 321-04 and 321-05 (**Fig S6D)**.

### The trajectory of TFH maturation in Flu-specific clonal lineages

We next sought to determine if the expanded and persistent TFH lineages we identified were in fact specific for Flu-derived antigens. Twelve paired TCRs were picked from donor 321-05 prioritized based on whether the lineage was detected in both PBMC and LN samples in the 10x dataset, the total number of cells detected in the lineage, and the number of time points the lineage was detected (**Fig 7A**, Extended Data Table 1). Four lineages (3, 9, 11, and 12) were detected in both the PBMC and LN samples 10X datasets, and for each clone the cTFH cells were detected either before or at the same time they were initially detected in the LN. Lineage 3 was detected in the cTFH at the time of initial vaccination on day 0 of year 1, indicating it is a memory response to Flu. Seven additional lineages (1, 2, 4, 5, 6, 7, and 10) were detected in bulk cTFH cells sequenced on day 5 (**Fig 7B**). Lineage 8 was not detected either in blood or LN prior to day 12 (**Fig 7A-B**), suggesting it is a *de novo* response. The paired TCRs were transduced into TCRnull Jurkat cells, and artificial antigen-presenting cell lines (aAPCs), here K562 cells, stably expressing each individual MHC class II allele from donor 321-05 were generated. First, we set out to determine the Flu protein containing the epitope recognized by each TFH clone by co-culturing them with aAPCs either a) infected with Flu PR8 strain (the vaccine backbone) and b) transfected with plasmids expressing individual segments of Flu PR8 strain and examining CD69 upregulation, IFNγ production, and CD3 downregulation as measures of activation **(Fig S7A).** Clones TFH1, TFH3, and TFH12 were responsive to PR8-infected aAPCs expressing DR3*03:01, DPA1*01:03 / DPB1*13:01, and DR5*01:01, respectively **(Fig S7B)**. Clone TFH1 responded to the matrix (M) gene segment with DR3*03:01 aAPCs, TFH3 also recognized M protein but in the context of DPA1*01:03 / DPB1*13:01, and TFH12 responded to the nucleoprotein segment (NP) in DR5*01:01 aAPCs **(Fig S7C)**. We synthesized partially overlapping 17mer and 15mer peptides tiled across the span of the M and NP gene segments, respectively, to precisely map the epitopes recognized by clones TFH1, TFH3, and TFH12. Each clone responded to two partially overlapping peptides from the same gene; TFH1 responded best to M2_46-62_, TFH3 to M1_51-67_, and TFH12 the NP_30-43_ (**Fig 7C**). We further tested if any of the remaining clones were specific for peptides derived from the HA protein by co-culturing with the T cell lines with aAPCs pulsed with full-length recombinant protein for each Flu strain included in the vaccine. Clone TFH11 showed a positive response to the HA from the H1N1 strain presented by DR5*01:01 aAPCs **(Fig. S7D)**. To more precisely map the location of the epitope recognized by TFH11, we tested whether it responded to aAPCs transfected with plasmids containing full-length and truncated versions of HA. Truncations were made by removing blocks of 100 amino acids from the C terminus of HA. TFH11 responded to constructs containing the first 300 amino acids of HA starting from the N-terminus but failed to respond once HA_160-260_ was deleted, indicating its epitope is located within this region of HA (**Fig. 7D).**

**Figure 7.**
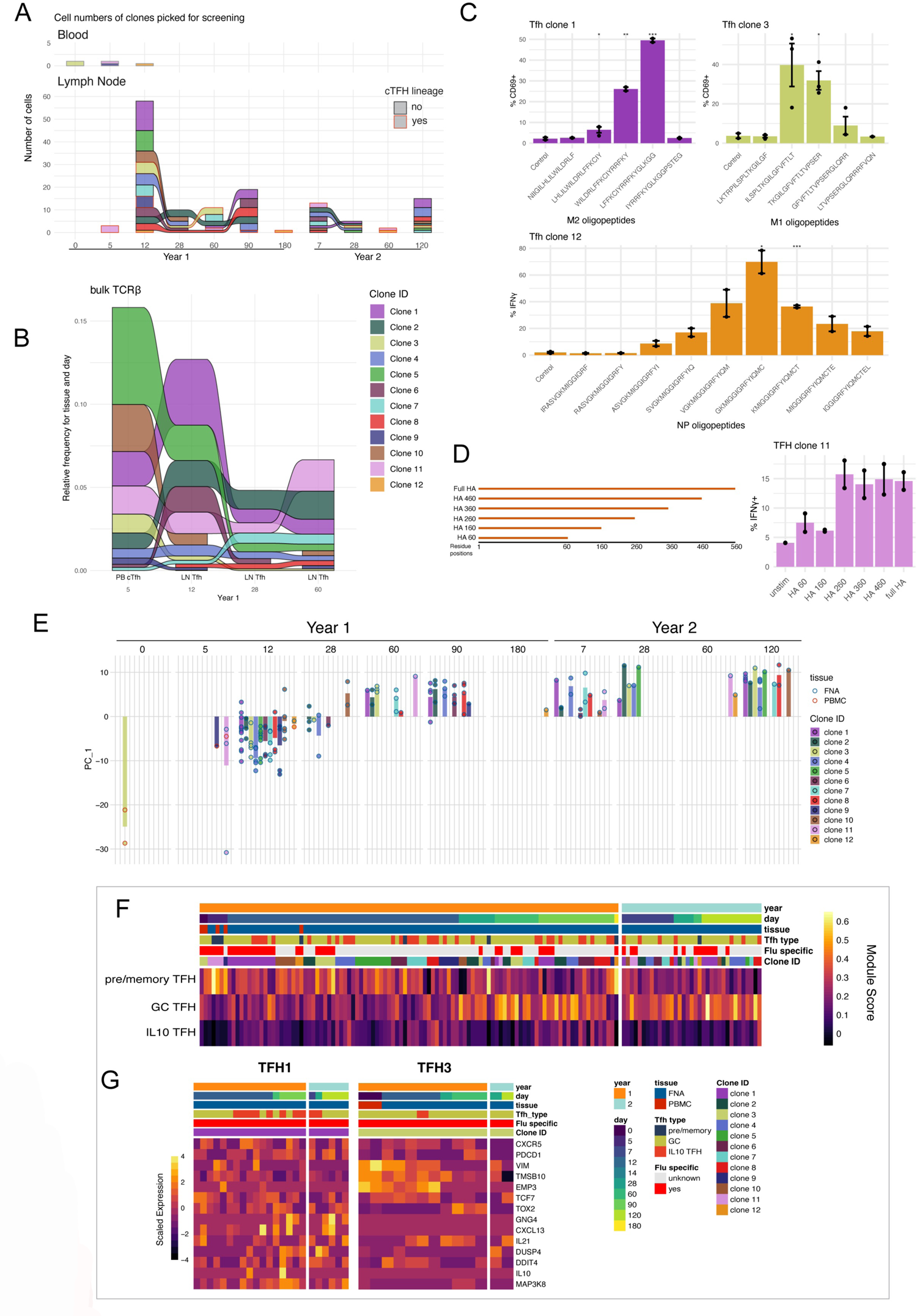
Dynamic alteration of phenotypes in Flu-specific TFH clonal lineages. **A)** Alluvial plot showing the number of cells detected in PBMCs (top) and LN (bottom) for each TFH clonal lineage picked from donor 321-05 for screening across time points. **B)** Alluvial plot showing the relative abundance of picked clonal lineages in bulk TCRβ sequencing from sorted cTFH and LN TFH cells. **C)** Frequency of CD69+ or IFNγ+ TFH1, TFH3, and TFH12 Jurkat T cells after co-culture with aAPCs pulsed with indicated IAV-derived peptides. **D)** Frequency of IFNγ+TFH12 Jurkat T cells after co-culture with aAPCs transfected with plasmids containing truncated versions of HA. **E)** PC1 scores of individual cells from the picked TFH lineages with respect to time. **F)** Heatmap of pre/memory, GC, and IL10 TFH gene set module scores for the picked TFH lineages. **G)** Heatmap of TFH marker gene expressions for TFH clones 1 and 9. In C and D the bar height corresponds to the mean, the whiskers show the standard error of the mean, and p-values calculated by t-test.

We next looked for alterations in GEX over the time course for all the screened clonal lineages to define their individual differentiation trajectories. Subsetting all cells belonging to each lineage and performing PCA analysis on their most highly variable features, we found the PC1 scores varied substantially within each lineage over the time course. The lowest positive PC1 scores were observed on days 0, 5, and 12 of year 1, ascended to values around 0 at day 28, and continued increasing until day 60 where they remained for the rest of year 1 and at all time points in year 2 (**Fig 7E**). We hypothesized this PCA analysis was capturing TFH maturation within these clonal lineages and more closely examined the expressions of genes contributing most to the variance for PC1, and indeed, the negative loadings contained a number of pre/memory TFH markers (e.g. *TMSB10, KLF2*) while the positive loadings were enriched with GC TFH markers (e.g. *SH2D1A*, *TOX2*) **(Fig S7E).** Scoring the expressions of gene sets specific to the pre/memory, GC, and IL10 TFH subsets as individual modules further confirmed the maturation of clonal lineages with the passage of time with the pre/memory module scores being highest early in year before decreasing as the GC and IL10 TFH scores increased with time (**Fig 7F**, Extended Data Table 2). Consistent with our time point analysis above, the pre/memory module scores remained low throughout the later part of year 1 and through year 2. Closer examination of TFH subset-specific genes within cells belonging to the Flu-specific TFH1 and TFH3 clonotypes found the daughter cells were heterogeneous with regard to TFH subset (**Fig 7G**). For example, the TFH1 and TFH3 lineages both contained a mix of mature IL10 and GC TFH cells across all time points. In the case of the TFH3 lineage where cells from the day 0 year 1 time point were detected there is a clear transition away from a migratory phenotype (i.e. high *VIM*, *TMSB10*, *EMP3* expression) at days 0 and 5 that is down regulated at later times presumably following migration into the follicle.

## Discussion

Our study describes the maturation of the TFH response to seasonal influenza vaccination using longitudinal samples taken from lymph nodes and peripheral blood from human donors over the course of two years.

TFH differentiation occurs on a continuum, however, we observed definable phenotypic states along this trajectory that correspond with their tissue location at both the macro and micro levels^1^. We show human TFH cells in the lymph node fall into three distinct subsets with definable transcriptional markers and differing metabolic programs. The pre/memory subset corresponded to TFH cells at the earliest and latest time points in the response when their frequency amongst TFH cells was highest; early during the beginning stages of differentiation at the interfollicular zone after initial antigen stimulation, and later after disengaging from the GC reaction and forming memory. In GC TFH cells we observed downregulation in the expression molecules related to cell motility and upregulation of *CXCL13* and *IL21*, two key mediators for B cell migration, retention, and survival within the GC. Increased glucose metabolism through mTOR signaling is required for differentiation into GC TFH cells in mouse models ^23^. In agreement with this, we observed upregulation of gene modules related to mitochondrial respiration and glycolysis in human GC and IL-10+ TFH cells compared to pre/memory and subsets, indicating this requirement is conserved between species. The existence of human FOXP3-IL-10+ TFH cells with regulatory functions were recently reported ^15,16^. In addition to sharing a core gene expression profile with GC TFH cells, IL-10+ TFH cells had several specific alterations that could hint at a potential mechanism for their differentiation, including suppression of *TCF7* expression associated with increased levels of *PRDM1*, specific expression of *FOXB1* (though no expression of *FOXP3)*, and modulated display of co-stimulatory/inhibitory molecules (e.g. increased *ICOS*, *LAG3*, and *GITR*).

While the existence of these three TFH subsets is known, the kinetics of TFH differentiation and the interrelatedness of these subsets within clonal lineages was unknown. Here, we show that within the LN TFH compartment that the pre/memory TFH subset had the highest frequency prior to vaccination, before decreasing over the course of several months as the frequency of GC TFH cells increased. Coinciding with the peak in GC TFH frequency was the appearance and increase in IL-10+ TFH cells. The frequency of IL-10+ TFH cells was low (< 5%) during the first month after vaccination, and only transiently increased and peaked in frequency between two and three months. Consistent with memory recall, TFH activation and expansion operated under different kinetics after vaccination in subsequent years. While it took three months for GC TFH cells to reach their peak frequency after vaccination during year one, they were the most abundant TFH subset by one week and reached their peak frequency within one month after re-vaccination during year two. The increased kinetics of GC TFH expansion and differentiation during the recall response in year two was accompanied by alterations in their transcriptional programs compared to the first year, including upregulated mitochondrial respiration and glycolytic pathways. Long-lived TFH cells in mice have been shown to undergo epigenetic alterations enforcing their metabolic reprogramming and supporting their survival and plasticity^24^. It is likely the transcriptional and metabolic alterations we observed in long-lived human TFH cells are regulated through similar epigenetic mechanisms and should be subject to further investigation.

Using TCR sequencing permitted us to track individual TFH clonotypes over the time course and examine for variation in their GEX profiles. Previous work comparing TFH cells from tonsils to cTFH in the blood for a single donor has shown these two compartments at least partially overlap in their TCR repertoires ^10,13^. Our analyses of bulk and single-cell TCR repertoires of LN TFH and cTFH revealed a similar overlap between the two compartments. Extending on this, we identified TFH clonotypes that were Flu-specific and persisted in the LN three to six months after initial vaccination in year one and reappeared in the same LN upon re-vaccination in year two where they again persisted for several months. By examining the GEX profiles of these Flu-specific TFH clonal lineages we were able to show that each TFH transcriptional state is attainable amongst daughter cells within a given lineage. The M1-specific TFH3 lineage was particularly interesting in that it was first detected as cTFH cells before vaccination on day 0 before appearing in the LN by day 12, indicating it was a memory response and that cTFH cells serve as a memory pool for seeding the GC reaction with antigen-specific TFH cells that assist B cells during the recall response.

It is notable that these robust persistent responses were observed with a vaccine that is unadjuvanted and generally considered to induce relatively weak immune responses. In contrast to other recent reports examining GC responses to mRNA vaccines or novel adjuvants, these responses induced expansion on a similar time scale (∼6 months), indicating that this may be a standard lifecycle of human TFH expansion, regardless of the inflammatory environment at priming ^14,25–27^. None of the subjects reported either an influenza infection or vaccination within the last three years, though this is not definitive evidence of a lack of exposure. However, the distinct dynamics between years 1 and 2 do suggest a resting environment at the year 1 time point.

In summary, our study shows the human TFH cells operate in a multi-tissue network that is plastic, dynamic, and durable across time. TFH cells provide critical help in the development of the humoral immune response to pathogen infection, vaccination, and autoimmunity. These works provide an extensive, high-resolution resource describing the evolution of the TFH response. Limitations of the study include the limited number of subjects and sampling, and our lack of knowledge of antigen exposure history. Future studies with expanded subjects and diverse vaccination types using the same antigen targets (e.g. adjuvanted vs. unadjuvanted Flu vaccination) will provide additional insights into the generation of productive TFH responses.

### Limitations of the study

A limitation of our study is that we examined a small number of individuals because of the cost and procedural intensity of the experimental plan. Future studies will build on these observed trajectories to characterize human TFH responses in more individuals and with diverse vaccination strategies.

## Supplemental Figure Legends

**Supplementary Figure 1.**
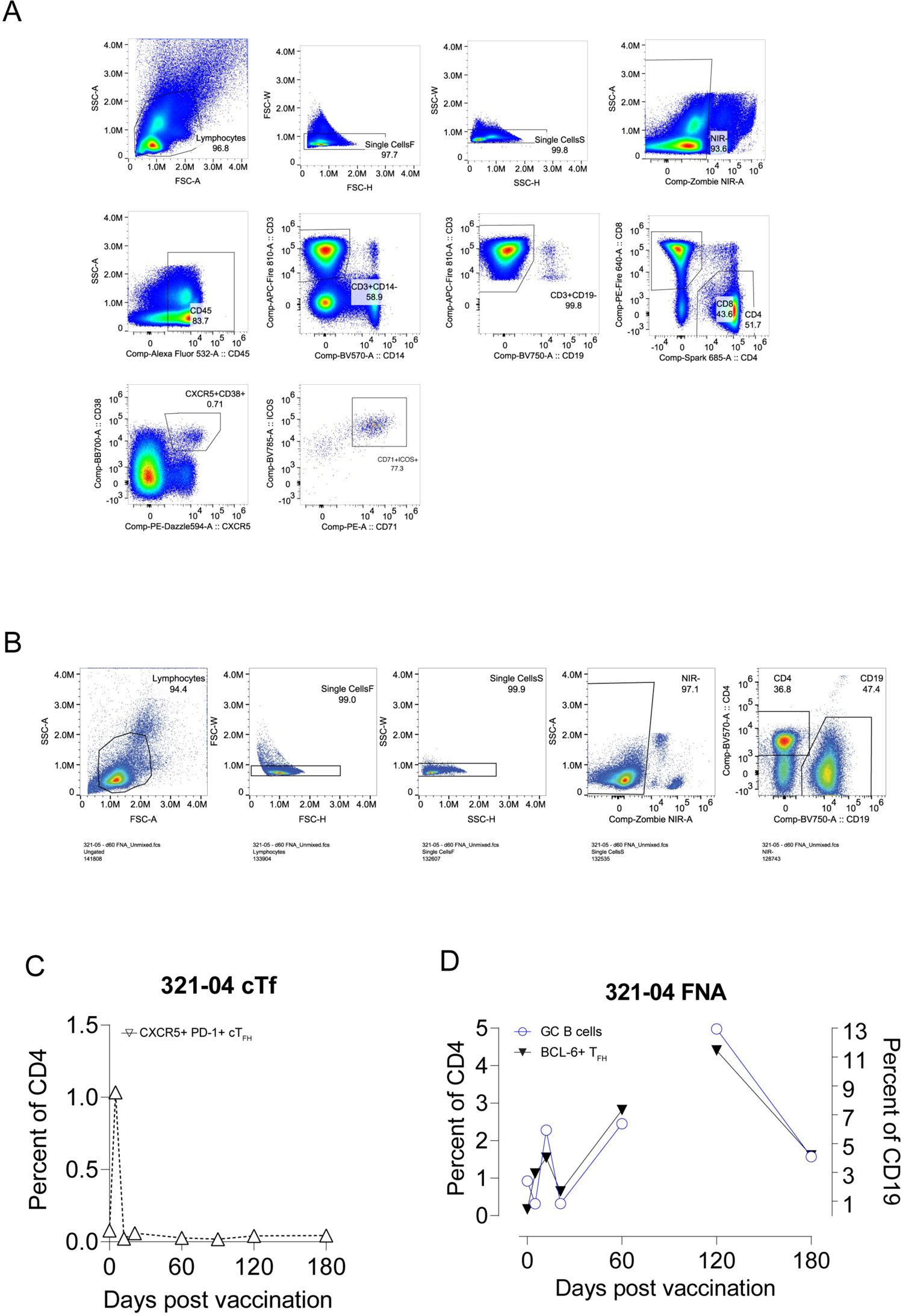
TFH flow gating strategy and donor 321-04 frequencies.

**Supplementary Figure 2.**
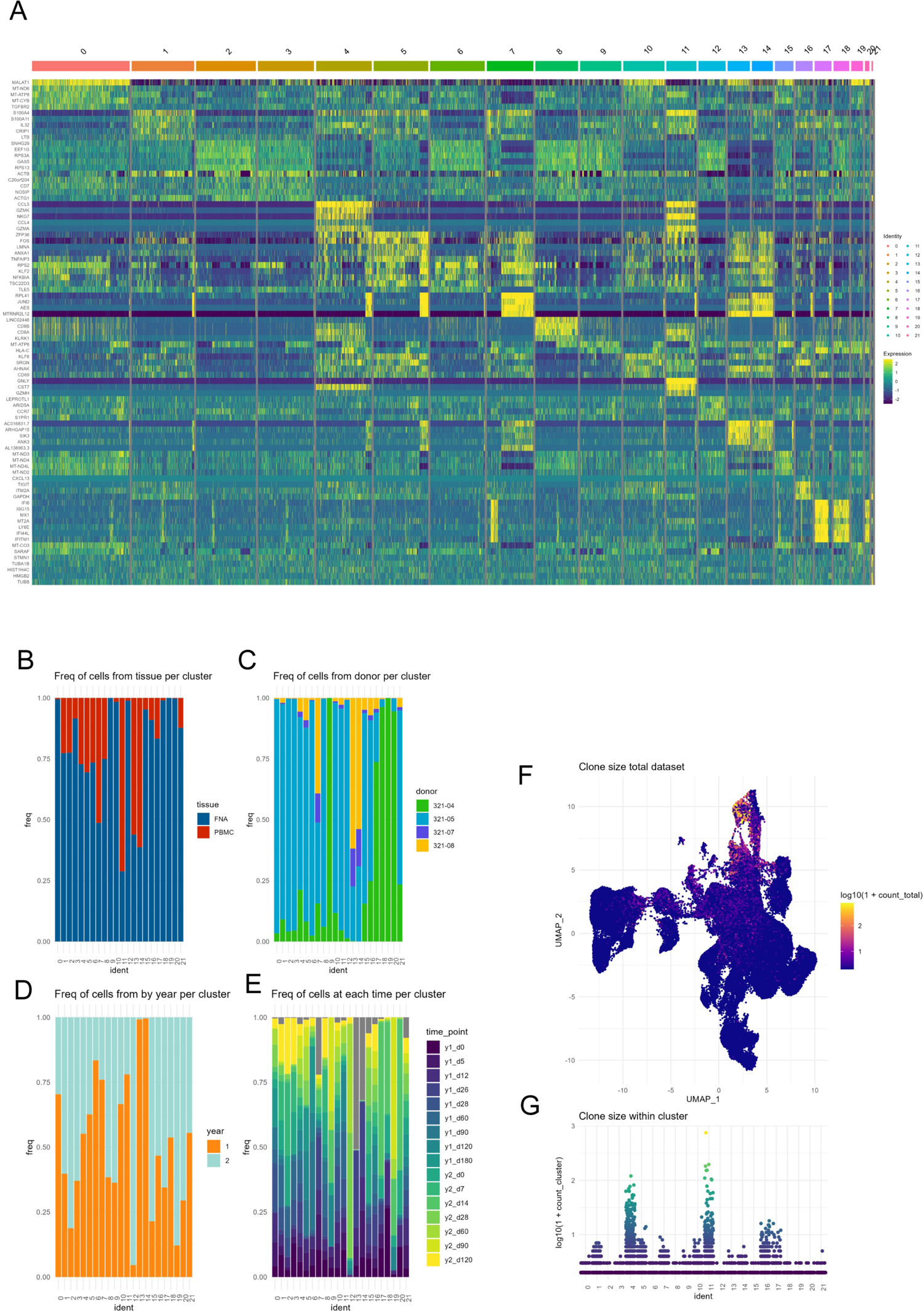
Features of the total T cell dataset. **A)** Heatmap showing the expressions of marker genes for each GEX cluster. **B-E)** GEX cluster distribution with respect to **B)** tissue, **C)** donor, **D)** study year, and **E)** time point after vaccinations. **F)** 2D umap projection colored by log10(1+n) transformed clone sizes. **G)** GEX cluster distribution of log10(1+n) transformed clone sizes.

**Supplementary Figure 3.**
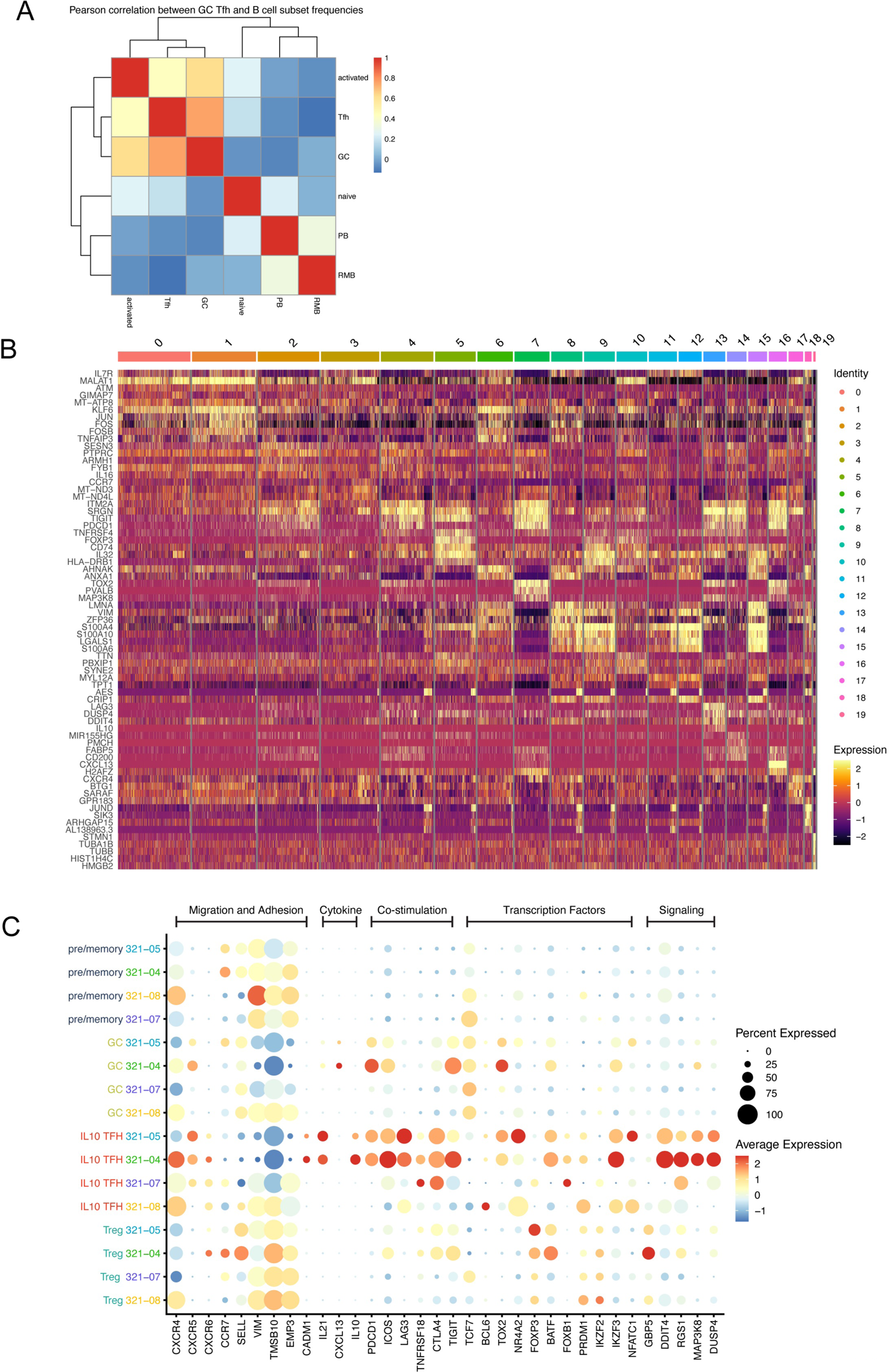
B cell subset frequency correlation and marker genes for TFH clonal lineages subset. **A)** Heatmap of Pearson correlation between frequency of TFH and naive, resting memory, activated, GC, and plasmablast (PB) B cell subsets in the lymph node. **B)** Heatmap of the top marker genes for each GEX cluster for the TFH clonal lineages subset. **C)** Dot plot of select TFH and Treg subset-specific markers split by donor.

**Supplementary Figure 4.**
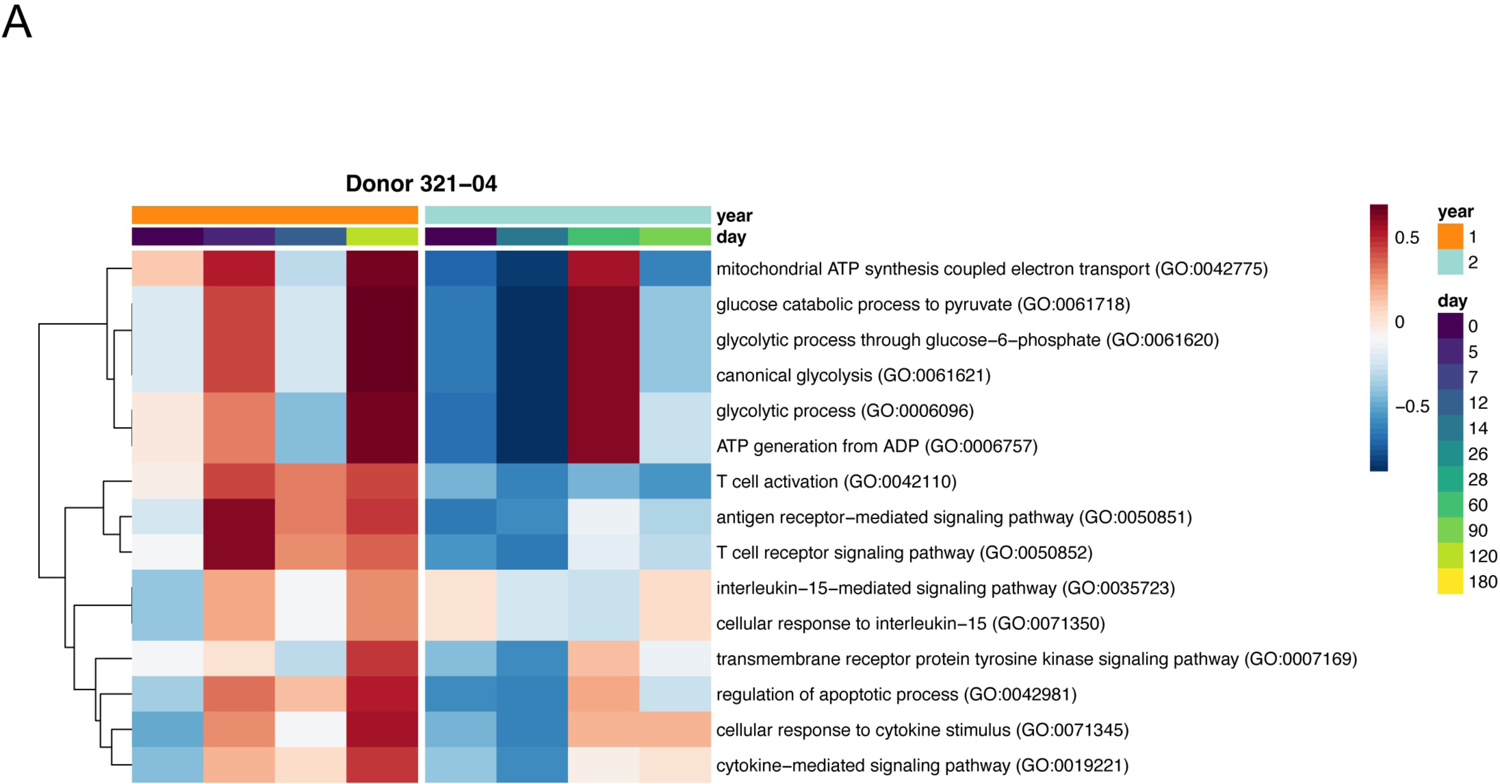
Alterations in TFH metabolism and signaling in Donor 321-04. **A)** GSVA of donor 321-05 LN TFH cells with respect to time for TFH-specific upregulated GO terms.

**Supplementary Figure 5.**
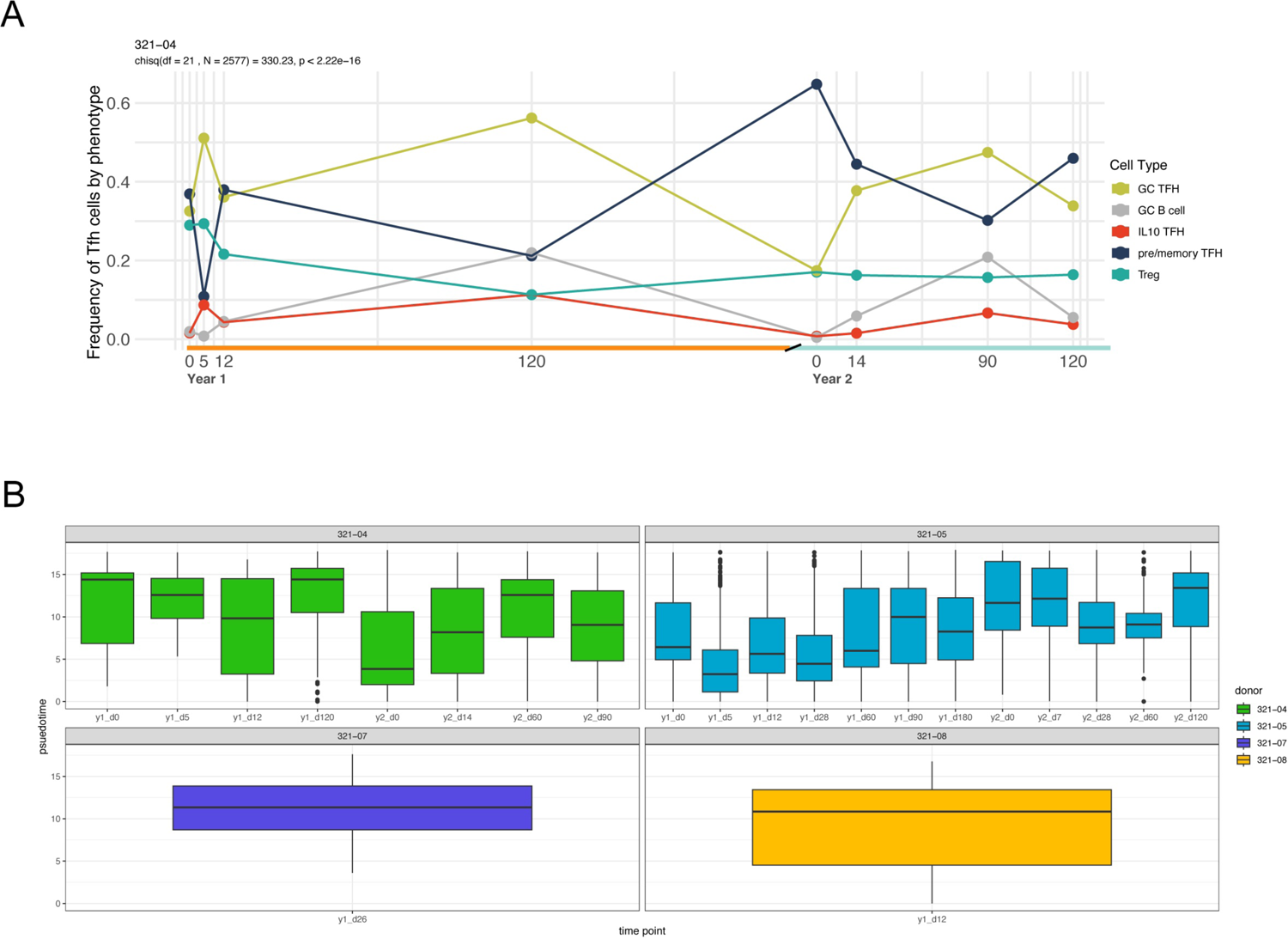
TFH composition and phenotypes over time for Donor 321-04. **A)** Relative frequencies of pre/memory, GC, and IL10 TFH subsets in LN samples from donor 321-04 over time. Chi-squared with p-value determined by Monte Carlo simulation. **B)** Boxplot of pseudotime values per cell for indicated donor and time points.

**Supplementary Figure 6.**
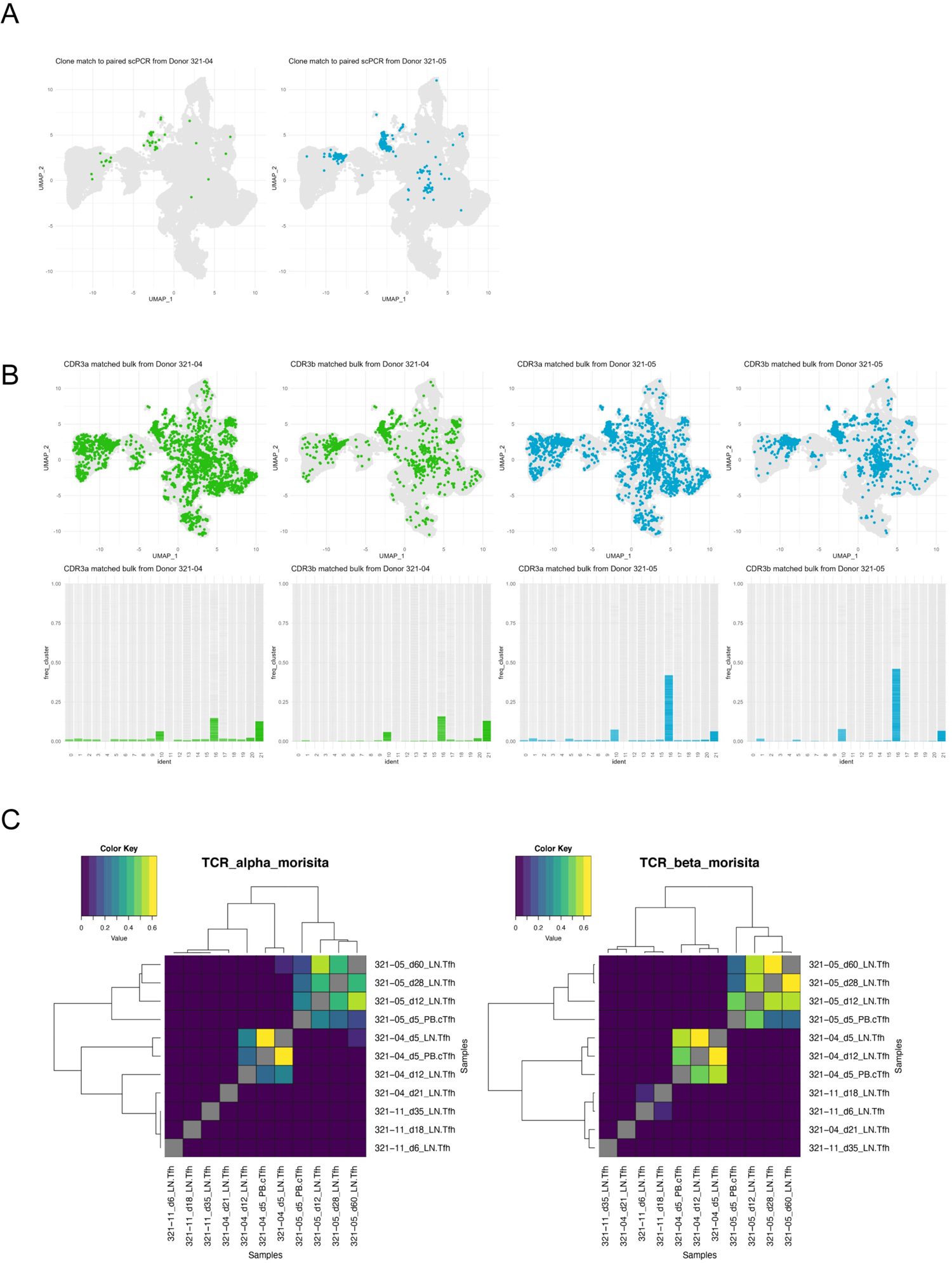
Bulk and single-cell PCR TFH repertoire profiling. **A)** 2D umap projection of total T cell dataset highlighting cells with matching paired TCR sequences to TCR data generated by scPCR of individually sorted cTFH and LN TFH cells from each donor. See Figure 1A. **B)** Top) 2D umap projection of total T cell dataset highlighting cells with matching TCRα or TCRβ sequences to TCR sequences generated from bulk sorted TFH cells from each donor. Bottom) Frequency cells within each GEX cluster matching to TCRα or TCRβ sequences in the bulk data. **C)** Morisita-Horn index for TCRα (left) and TCRβ (right) sequences from bulk sorted TFH cells for all donors and time points.

**Supplementary Figure 7.**
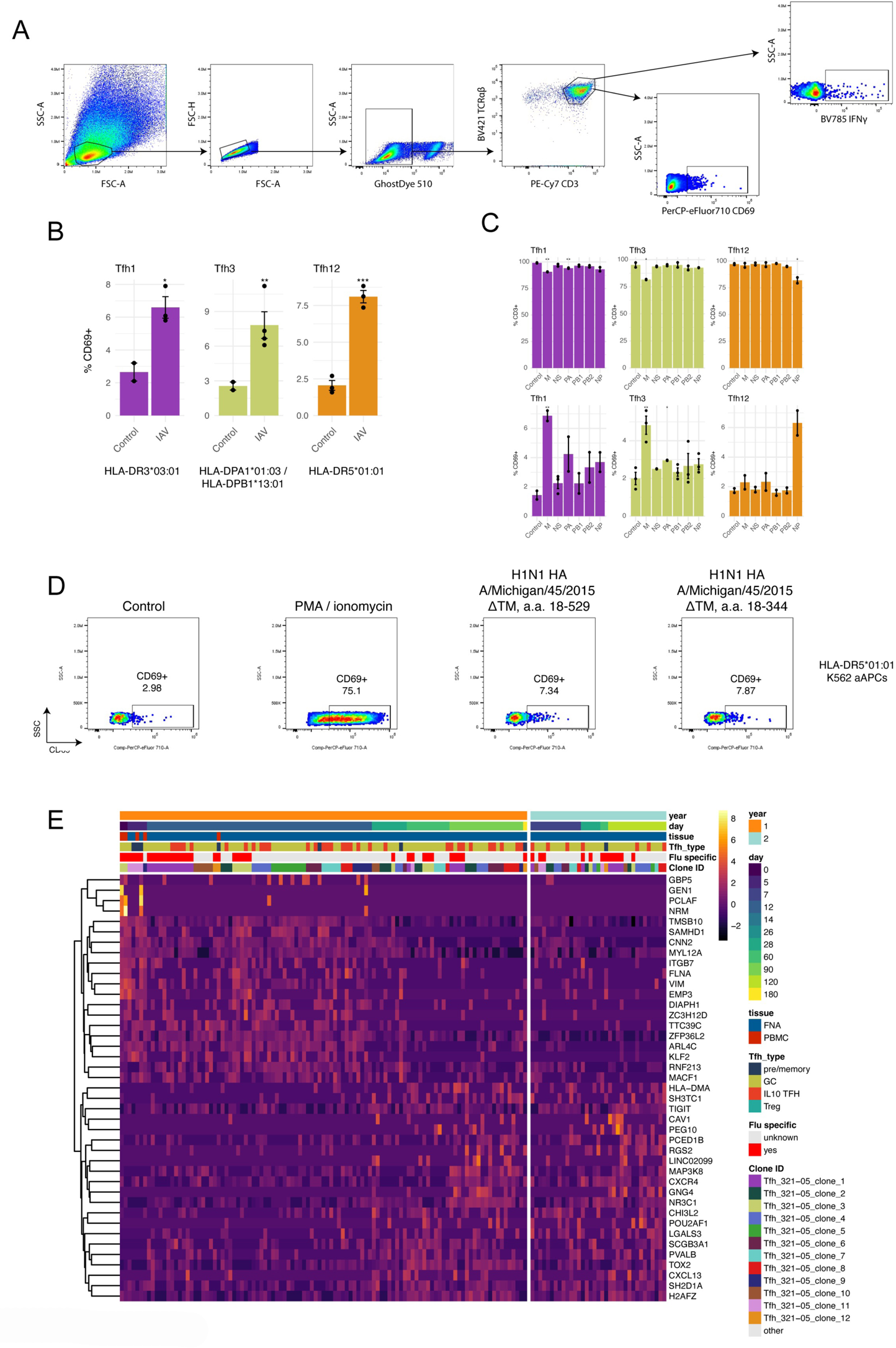
Identification of Flu-specific TFH clonotypes in donor 321-05. **A)** Gating strategy for assessing Jurkat TFH TCR cell line activation in co-culture experiments. **B)** Frequency of CD69+ Jurkat T cells expressing TCRs TFH1, TFH3, and TFH12 after co-culture with aAPCs infected with Flu PR8. **C)** Frequency of CD3+ (top) and CD69+ (bottom) TFH1, TFH3, and TFH12 T cell lines after co-culture with aAPCs transfected with plasmids expressing individual segments of the IAV genome. **D)** Frequency of CD69+ TFH11 cell line co-culture with aAPCs pulsed with recombinant HA protein, PMA/ionomycin, or unstimulated control. **D)** Heatmap showing the expressions of genes corresponding to the head and tail PC1 loadings in the picked TFH clonal lineages.

Extended Data

Extended Data Table 1. TFH TCRs chosen for specificity screening.

Extended Data Table 2. TFH subset module gene sets.

Extended Data Table 3. Study participant HLA typing.

## Methods

### Sample collection, preparation, and storage

All studies were approved by the Institutional Review Board of Washington University in St Louis. Written consent was obtained from all participants. Eight participants who had not been vaccinated against influenza for at least three years were enrolled, including 1 female and 7 males, aged 26–40 years old. PBMCs were isolated using Vacutainer CPT tubes (BD), the remaining red blood cells were lysed with ammonium chloride lysis buffer (Lonza), and cells were immediately used or cryopreserved in 10% dimethylsulfoxide in FBS. Ultrasound-guided FNA of axillary lymph nodes was performed by a qualified physician’s assistant under the supervision of a radiologist. Lymph node dimensions and cortical thickness were measured before each FNA. For each FNA sample, 6 passes were made using 25-gauge needles, each of which was flushed with 3 ml of RPMI 1640 supplemented with 10% FBS and 100 U ml−1 penicillin/streptomycin, followed by three 1-ml rinses. Red blood cells were lysed with ammonium chloride buffer (Lonza), washed twice with PBS supplemented with 2% FBS and 2 mM EDTA, and immediately used or cryopreserved in 10% DMSO in FBS. Participants reported no adverse effects of phlebotomy, serial FNA, or vaccination. No statistical methods were used to predetermine sample size. Investigators were not blinded to experiments and outcome assessment.

### Vaccine

Flucelvax QIV influenza vaccine for the North American 2018/2019 and 2019/2020 seasons were purchased from Seqirus.

### Single-cell RNA-seq library preparation and sequencing

Activated and memory B cells were enriched from PBMCs by first staining with IgD-PE and MojoSort anti-PE Nanobeads (BioLegend), and then processing with the EasySep Human B Cell Isolation Kit using the EasyEights magnet (Stemcell) to negatively enrich IgDlo B cells. Enriched IgDlo B cells, whole PBMCs, and whole FNA from each time point for participant 05 were processed using the following 10 × Genomics kits: Chromium Single Cell 5′ Library and Gel Bead Kit v2 (PN-1000006); Chromium Single Cell A Chip Kit (PN-120236); Chromium Single Cell V(D)J Enrichment Kit; and Human, B cell (96rxns) (PN-1000016), and Chromium i7 Multiplex Kit (PN-120262). The cDNAs were prepared after GEM generation and barcoding, followed by GEM RT reaction and bead cleanup steps. Purified cDNA was amplified for 10–14 cycles before cleaning with SPRIselect beads. Then, samples were evaluated on a bioanalyser to determine cDNA concentration. BCR target enrichments were performed on full-length cDNA. GEX and enriched BCR libraries were prepared as recommended by the 10 × Genomics Chromium Single Cell V(D)J Reagent Kit (v1 Chemistry) user guide, with appropriate modifications to the PCR cycles based on the calculated cDNA concentration. The cDNA libraries were sequenced on Novaseq S4 (Illumina), targeting a median sequencing depth of 50,000 and 5,000 read pairs per cell for gene expression and BCR libraries, respectively

### Cell sorting and flow cytometry

Staining for analysis and sorting was performed using fresh or cryo-preserved PBMCs or FNA single cell suspensions in 2% FBS and 2 mM EDTA in PBS (P2). For sorting, cells were stained for 30 min on ice with IgD-PerCP-Cy5.5 (IA6-2, 1:200), CD4-Alexa 700 (SK3, 1:400), CD20-APC-Fire750 (2H7, 1:100), and Zombie Aqua along with CD38-BV605 (HIT2, 1:100), CD71-FITC (CY1G4, 1:200), and CD19-PE (HIB19, 1:200) for PBs or CD19-BV421 (HIB19, 1:100), CD71-PE (CY1G4, 1:400), CXCR5-PE-Dazzle 594 (J252D4, 1:40), and CD38-PE-Cy7 (HIT2, 1:200) for GC B cells (all BioLegend). For donors 321-07 and 321-08 the PBMC samples for days 0 and 5 were stained with TotalSeq-C anti-human hashtag oligos 9 and 10, respectively, for downstream demultiplexing. Cells were washed twice, and single PBs (live singlet CD19+ CD4− IgDlo CD38+ CD20− CD71+) and GC B cells (live singlet CD19+ CD4− IgDlo CD71+CD38int CD20+ CXCR5+) were sorted using a FACSAria II into 96-well plates containing 2 μL Lysis Buffer (Clontech) supplemented with 1 U μl−1 RNase inhibitor (NEB), or bulk sorted into buffer RLT Plus (Qiagen) and immediately frozen on dry ice. For analysis, cells were stained for 30 min on ice with biotinylated recombinant HAs and PD-1-BB515 (EH12.1, BD Horizon, 1:100) diluted in P2, washed twice, then stained for 30 min on ice with IgA-FITC (M24A, Millipore, 1:500), CD45-PerCP (2D1, BD Bioscience, 1:25), IgG-BV480 (goat polyclonal, Jackson ImmunoResearch, 1:100), IgD-SB702 (IA6-2, Thermo, 1:50), CD38-BV421 (HIT2, 1:100), CD20-Pacific Blue (2H7, 1:400), CD27-BV510 (O323, 1:50), CD4-BV570 (OKT4, 1:50), CD24-BV605 (ML5, 1:100), streptavidin-BV650, CD19-BV750 (HIB19, 1:100), CXCR5-PE-Dazzle 594 (J252D4, 1:50), CD71-APC (CY1G4, 1:100), CD14-A700 (HCD14, 1:200), and IgM-APC-Cy7 (MHM-88, 1:400) (all BioLegend) diluted in Brilliant Staining buffer (BD Horizon). Cells were washed twice, then fixed and permeabilized for intranuclear staining for 1 h at 25 °C with True Nuclear fixation buffer (BioLegend), washed twice with permeabilization/wash buffer, and stained for 30 min at 25 °C with BCL6-PE (7D1, 1:50) and Ki-67-PE-Cy7 (Ki-67, 1:400) (both BioLegend). Cells were washed twice with permeabilization/wash buffer and resuspended in P2 for acquisition on an Aurora using SpectroFlo v2.2 (Cytek). Flow cytometry data were analysed using FlowJo v10 (Treestar).”

### Processing of 10× Genomics single-cell 5′ gene expression data

The demultiplexed FASTQ pair-end reads were preprocessed on a by-sample basis using the `cellranger count` command from 10× Genomics’ Cell Ranger v6.0.0 for alignment against the GRCh38 human reference v.3.0.0 (refdata-cellranger-GRCh38-3.0.0). Individual samples are assigned to one of three datasets: one containing samples from donors 321-05 and 321-04, one containing samples from donors 321-07 and 321-08, and the last containing additional CD4+ PBMCs from donor 321-05 year 1 day 0 time point.

### Processing 10× Genomics single-cell TCR reads

Read for each sample were parsed using `cellranger vdj` from 10 × Genomics’ Cell Ranger v.6.0.0 for alignment against the cellranger-vdj-GRCh38-alts-ensembl-5.0.0 reference. Additional quality control was performed using the `make_10x_clones_file` function from the `CoNGA` software package using settings `stringent = True` to correct for spurious chain pairings based on other paired clonotypes in the dataset. Only cells with paired TCRαβ sequences were retained. The inverse D50 index in figure 6D was calculated as the reciprocal of the quotient from dividing the number of unique clones occupying the top 50% of the repertoire (after ranking by abundance) by the total number of unique clonotypes for the time point multiplied by 100.

### Processing single-cell TCR sequencing from sorted TFH cells

TCRα and TCRβ chains were amplified from blood and lymph node TFH cells sorted into individual wells of a 384-well plate (described above) by RT-PCR using variable gene and constant region primers and sequenced by Sanger method as previously described ^28^. Sequencing reads were parsed using the TCRdist pipeline as previously described ^29^.

### Bulk TCR sequencing of sorted TFH cells

TCRα and TCRβ chains from bulk sorted cTFH and LN TFH were amplified using a 5’ Rapid Amplification of cDNA Ends (RACE) with unique molecular identifiers (UMIs) for error correction essentially as described^30^. RNA was extracted from bulk sorted blood and lymph node TFH cells from donors 321-04, 321-05, and 321-11 using the RNeasy Micro Kit (Qiagen). Reverse transcription was carried out using SmartScribe RT (Takara), and Q5 polymerase (New England Biolabs) was used during first and second round amplification. Barcoded TCRα and TCRβ amplicons generated by the second round PCR were pooled by equal volume, prepped, and indexed for sequencing on Illumina platforms using a KAPA HyperPrep Kit (Roche). 150 bp paired-end sequencing was performed on Illumina NovaSeq6000.

### Processing bulk TCR sequencing

Paired-end FASTQ reads were processed using the `migec v1.2.9` software ^31^. Reads corresponding to individual samples were demultiplexed prior to assembling nucleotide sequence reads within each UMI group. VDJ junction mapping of assembled contigs was performed using the `NCBI-BLAST+` package within `migec.` Additional quality control was performed on the filtered clonotype tables from `migec` using the `vdjtools v1.2.1` ^32^ `FilterNonFunctional`, `Correct`, and `Decontaminate` functions to remove potentially erroneous clonotypes within and between samples. The `immunarch v0.6.5` package ^33^ for R was used for the Morisita-Horn index calculations.

### Single-cell gene expression analysis

Analysis was performed using `Seurat v 4.1.1`. The three datsets noted above were each processed in the follwing manner prior to aggregation. First, an object containing all cell types was generated prior to filtering down to only T cells. QC was performed by removing cells with greater than 10% mitochondrial content, expressing fewer than 300 or greater than 4000 genes, and containing fewer than 900 or greater than 40000 UMIs were filtered from the dataset to remove presumably lysed/dead cells and doublets. `MULTIseqDemux` for demultiplexing hashtag oligos for the datset containing donors 321-07 and 321-08 ^34^. After preprocessing, the three datasets were merged and `SCTranform` was used for data normalization prior to feature extraction, scaling, and principal component analysis (PCA) ^35^. The `Harmony` algorithim was applied for batch integration across donors and datasets ^36^. The `FindClusters` and `RunUMAP` tools were applied to the harmonized embedding generate a 2D projection of GEX space. TCR clonotype information (described above) was then mapped back to cells in the object by matching their corresponding UMI barcodes.

Cluster identities were assigned based on the expression of T cell markers distinguishing each of the major subtypes including: *CD4*, *CD8B*, *KLRB1*, *ZBTB16*, *KLRG1*, *CCR7*, *SELL*, *FOXP3*, *PRF1*, and *TBX21*. *PDCD1*, *CXCR5*, and *ICOS* were used to identify TFH clusters. TFH cells were further subset on and reprocessed to identify heterogeneity in their GEX phenotypes. Here, we chose to include all cells clonally-linked to those in the two TFH clusters that might otherwise be excluded due to non-TFH GEX cluster assignments.

Monocle3 ^37–39^ R package version 1.1.0 was used for pseudotime analysis on our data. The `getEnrichedPathways` function in the Cerebro ^40^ v1.3.1 R package was used to find enriched GO terms for the TFH populations. GO terms enriched in IL10 TFH cells with respect to all other TFH and Treg cells were used for GSVA ^41^ analysis in Figure 4B (GSVA R package v1.42.0).

Genes defining the pre/memory, GC, and IL10 TFH modules in Figure 7F were taken from intersecting positive markers defining the subset relative to the two other subsets based on the results of the differential expression analysis shown in Figure 3G. Module scores were calculated using the `AddModuleScore` function in Seurat.

### HLA class II typing

HLA typing was conducted with the research-grade AllType NGS 11-loci Amplification Kit (One Lambda) as described in (https://doi.org/10.1038/s41590-022-01184-4). Libraries were sequenced on the Illumina MiSeq platform at 150×150 base pairs, and data were parsed using the TypeStream Visual software (One Lambda). HLA typing results are in Extended Data Table 3.

### Construction of TFH TCR cell lines and aAPC lineages

Twelve TCR clonotypes were picked from donor 321-05 for cloning based on two criteria; the clone was detected at multiple time points after vaccination, and it had matching scPCR and bulk sequencing results from the sorted TFH cells. VDJ nucleotide sequences of the picked clones were reconstructed using `stiTChR`^42^ with the TCRα and TCRβ chains linked with a P2A auto-cleavage site. To generate APCs expressing specific HLA, the nucleotide sequences of alpha and beta chains for donor-specific HLA class II haplotypes were curated using (<u>Allele</u> <u>Search Tool < IMGT/HLA < IPD < EMBL-EBI</u>) and linked via T2A site to form the full HLA constructs (HLA-DR, HLA-DP, and HLA-DQ). Constructs were synthesized and cloned (Genscript) into a custom pLVX-EF1a-P2A-GFP-IRES-Puro lentiviral vector. Lentivirus particles were subsequently generated by co-transfecting 293T cells (ATCC) with one of the generated constructs, psPAX2 (Addgene, 12260), and pMD2.G (Addgene, 12259) using GeneJuice transfection reagent (EMD Millipore) or Lipofectamine3000 transfection kit (Thermo Fisher). To generate the a-APC lineages, K562 cells (ATCC, CCL-243) expressing (HLA-DM, CD80, and CD64 molecules) were transduced with donor’s HLA-II vectors to generate a set of artificial Antigen Presenting Cell lines (aAPCs). Similarly, Jurkat cells were transduced with the panel TCR lentiviruses. Both K562 and Jurkat cells were placed under 1 ug/mL puromycin selection for 2 weeks before sorting for GFP+ cells.

### Cell line antigen stimulation assays

TCR-negative Jurkat cells were cloned with a curated set of donor’s derived TCR sequences. The TCR expressing clonotypes were subsequently screened for activation upon coculture with aAPC encoding cognate HLA Class-II alleles. The HLA-II-expressing aAPCs were preincubated with B-Propiolactone-inactivated Influenza A virus (strain A/Puerto Rico/8/1934 H1N1) at MOI=1 for 2 hours prior to adding the TCR expressing lines in a ratio of 1:1. The cocultured cells were incubated at 37°C and 5% C02 for additional 24 hours. Cells were washed with cold PBS and then stained with a cocktail of fluorescently labeled antibodies including: anti-human CD3-PE-Cy7 (Clone: OKT3, Biolegend), Ghost viability dye (Tonbo Biosciences),TCR a/b-BV421 (Clone: IP26, Biolegend), CD69-PerCP-eFluor-710(Clone: FN50, Invitrogen), IFN-g-BV785 (Clone: 4S.B3, Biolegend). Similarly, HLA-II expressing aAPCs were transfected with pHW2000 vectors encoding the genomic viral segments (PB2, PB1, PA, NP, M, and NS) 48h prior to coculturing with the TCR expressing lineages. The transfection was conducted via Neon Electroporation System (Thermo, MPK5000S) using 1 pulse at 1,000 volts and 50 ms width according to the manufacturer’s protocol. To identify the driving peptide motifs that triggered activation in the responding clones, a pool of overlapping oligopeptide sequences spanning M1, M2, and NP proteins of the H1N1 PR8 strain (Mimotopes) were used for the peptide mapping experiments. Libraries covering M1 and M2 consisted of 17mer peptides that were overlapping by 11 residues.The NP library consisted of 15mer peptides overlapping by 11 residue. All samples were acquired on Aurora (CyteK) and data were analyzed using FlowJo software.

## Acknowledgments

We thank Erica Lantelme for facilitating sorting; Lisa Kessels, Michael Royal, and the staff of the Infectious Diseases Clinical Research Unit at Washington University School of Medicine for assistance with vaccination and sample collection.

The authors declare the following competing interests: A.H.E. is a consultant for InBios and Fimbrion Therapeutics. The Ellebedy laboratory received funding under sponsored research agreements from Emergent BioSolutions. P.G.T. has consulted and/or received honoraria and travel support from Illumina, Johnson and Johnson, and 10X Genomics. P.G.T. serves on the Scientific Advisory Board of Immunoscape and Cytoagents. The authors have applied for patents covering some aspects of these studies. All other authors declare no competing interests.

The Thomas laboratory was funded by ALSAC at St. Jude; the Center for Influenza Vaccine Research for High-Risk Populations (CIVR-HRP) contract number 75N93019C00052; the St. Jude Center of Excellence for Influenza Research and Surveillance (P.G.T.) contract number HHSN272201400006C; the St. Jude Center of Excellence for Influenza Research and Response (P.G.T.) contract number 75N93021C00016; and grants U01AI150747, U01AI144616, and R01AI136514. The Ellebedy laboratory was supported by NIAID grants R21 AI139813, U01 AI141990, and NIAID Centers of Excellence for Influenza Research and Surveillance (CEIRS) contract HHSN272201400006C.

The WU321 study was reviewed and approved by the Washington University Institutional Review Board (approval no. 201808171).

## Contributions

S.A.S analyzed scGEX and TCR sequencing data, designed experiments, generated libraries and processed data for bulk TCR profiling, and wrote the manuscript. J.S.T. collected and analyzed the flow cytometry data and performed fluorescence-activated cell sorting. M.G. generated TCR and aAPC cell lines and performed antigen-stimulation experiments. J.C.C. and H.K. parsed and analyzed scGEX and TCR data. W.A. performed single-cell TCR sequencing. A.H.E., R.M.P., M.K.K. and A.H. wrote and maintained the IRB protocol, recruited, vaccinated and phlebotomized participants and coordinated sample collection. T.S. performed the FNA under the supervision of S.T., J.Q.Z., J.H. A.H.E. conceived, designed, and supervised the seasonal Flu vaccination study. P.G.T. supervised this study and wrote the manuscript. All authors reviewed the manuscript.

## Data Availability

Single-cell GEX and TCR profiling for all the T cells with paired TCR information along with the bulk and single-cell TCR sequencing of sorted TFH cells is available on Zenodo https://doi.org/10.5281/zenodo.6476022. The code for running the analyses is available at https://github.com/sschattgen/Flu_TFH_paper.

